# EndophilinA-dependent coupling between activity-dependent calcium influx and synaptic autophagy is disrupted by a Parkinson-risk mutation

**DOI:** 10.1101/2022.04.29.490010

**Authors:** Adekunle T. Bademosi, Marianna Decet, Sabine Kuenen, Carles Calatayud, Jef Swerts, Sandra F Gallego, Nils Schoovaerts, Nikolaos Louros, Ella Martin, Spyridoula Karamanou, Jean-Baptiste Sibarita, Katlijn Vints, Natalia V. Gounko, Frédéric A. Meunier, Anastassios Economou, Wim Versées, Frederic Rousseau, Joost Schymkowitz, Sandra-F. Soukup, Patrik Verstreken

## Abstract

Neuronal activity and neurotransmitter release cause use-dependent decline in protein function. However, it is unclear how this is coupled to local protein turnover and quality control mechanisms. Here we show that the endocytic protein Endophilin-A (EndoA/ENDOA1) couples activity-induced calcium influx to synaptic autophagy and neuronal survival. We identify single mutations in the EndoA flexible region that either increases EndoA diffusion and promotes autophagosome formation in the absence of calcium, or immobilizes EndoA and blocks autophagy, even in the presence of calcium. Hence, the EndoA flexible region is a switch that responds to calcium, regulating EndoA nanoscale synaptic organization and association with autophagosomes driving their formation. Interestingly, a pathogenic variant in the human ENDOA1 variable region that confers risk to Parkinson’s disease (PD), also confines ENDOA1 to the synaptic plasma membrane and equally blocks autophagy in flies *in vivo* and in induced human neurons. Thus, our work reveals a mechanism neurons use to connect neuronal activity to local protein turnover by autophagy, which is critical for neuronal survival.

## Introduction

Presynaptic terminals are complex machines that drive a multitude of functions such as memory acquisition, complex coordinated movements and thought (Mayford et al., 2012). Synapses are densely packed with proteins and lipids (Wilhelm et al., 2014) that power essential processes such as neurotransmitter release, changes in plasticity, endocytosis, signaling, etc. (Sudhof, 2004; Südhof, 2013; Südhof and Malenka, 2008). Yet synapses are often located far from neuronal cell bodies and thus, nerve terminals need to, at least in part, locally cope with turnover of bio-molecules (Azarnia Tehran et al., 2018; Decet and Verstreken, 2021; Soukup et al., 2018; Vijayan and Verstreken, 2017; Wang et al., 2017).

There are several cellular “homeostasis mechanisms”, that are involved in quality control and protein turnover at the synapses, including macroautophagy, where double membrane structures engulf parts of the cytoplasm destined for degradation and recycling. However, synapses are fragile and it is likely they avoid “blunt” and “massive” turnover by gorging sizeable parts of their cytoplasm. For example, based on electron microscopy studies, autophagosomes can measure up to hundreds of nanometers in diameter and thus a single autophagosome could engulf up to >10-20% of the synaptic cytoplasm at once (Baba et al., 1997; Jin and Klionsky, 2014; Klionsky et al., 2021; Soukup et al., 2016; Vanhauwaert et al., 2017). Hence, it is conceivable that autophagy at synapses is a well-regulated process.

There are several synapse-specific proteins that are implicated in the creation of autophagosomes at nerve terminals and these same proteins are not involved in autophagy elsewhere in the cell (Azarnia Tehran et al., 2018; Decet and Verstreken, 2021; Montenegro-Venegas et al., 2020; Soukup et al., 2016; Vanhauwaert et al., 2017). This includes the active zone protein Bassoon that binds to and sequesters Atg5, thus limiting the ability to create new autophagosomes at synapses (Okerlund et al., 2017). We have also implicated Synaptojanin-1 that dephosphorylates phosphoinositides at the autophagosomal membrane to facilitate Atg18-cycling (Soukup et al., 2018; Vanhauwaert et al., 2017). Similarly, we found a role for EndoA1, a small BAR and SH3 domain-containing protein that, when phosphorylated by LRRK2, creates membrane docking sites for autophagic proteins such as Atg3 or Atg1 (Murdoch et al., 2016; Soukup and Verstreken, 2017; Soukup et al., 2018, 2016). While the important question of how key molecular pathways trigger autophagy at synapses is not answered, these discoveries are starting to reveal exciting connections to neurodegenerative disease that we do not yet fully understand. Indeed, the genes encoding Synaptojanin-1 and LRRK2 have been found mutated in PD. There is also a GWAS signal close to the *LRRK2* locus and two independent GWAS signals in the vicinity of the *SH3GL2* gene that encodes ENDOA1 (Daida et al., 2020; Satake et al., 2009; Simón-Sánchez et al., 2009). Interestingly, there is also a mutation in the flexible region of ENDOA1, between the BAR and SH3 domains that confers increased risk to develop PD (Germer et al., 2019). However, the effect of this mutation on EndoA function is unknown.

Synapses require intense metabolic activity to power the vesicle cycle and other membrane-bound processes. This also causes protein stress and damage. An interesting idea is that this stress and damage are coupled to protein- and organelle-turnover including that, across species, neuronal activity induces the formation of autophagosomes at synapses (Decet and Verstreken, 2021; Hill et al., 2019; Kroemer et al., 2010; Kulkarni et al., 2021; Nakatogawa, 2020; Nixon, 2013; Shehata et al., 2012; Soukup et al., 2016; Wang et al., 2015; Yang et al., 2022). However, how elements of neuronal activity such as calcium influx, neurotransmitter release or endocytosis, drive the formation of autophagosomes at synaptic terminals has not been investigated.

Here, we show that neuronal stimulation-induced calcium influx is necessary and sufficient to drive the formation of autophagosomes at pre-synaptic terminals. This process is mediated by ENDOA1 and blocked by the pathogenic risk variant mutation, G276V, in the flexible region of the human protein. This region controls EndoA nanoscale organization at synapses in a calcium-dependent manner, such that at rest the protein is in the periphery, where it can promote synaptic vesicle endocytosis, while during stimulation, it relocalizes to the synapse lumen to facilitate autophagosome formation. Finally, balanced EndoA-dependent and stimulation driven synaptic autophagy is required for neuronal survival. Our work reveals the impact of the ENDOA1 risk variant and suggests a critical function for synaptic autophagy in PD.

## Results

### EndoA is required for Ca^2+^-induced pre-synaptic autophagy

It is not known how neuronal activity induces autophagy at synapses (Decet and Verstreken, 2021; Shehata et al., 2012; Soukup et al., 2016; Wang et al., 2015). We therefore tested if Ca^2+^ influx induced by stimulation can trigger the process. We performed live confocal imaging of *Drosophila* third-instar larval neuromuscular junction (NMJ) boutons expressing Atg8 fused to mCherry (Atg8^mCherry^) under endogenous *atg8-*promotor control. Motor neurons were electrically stimulated within the range of their normal physiological firing ability (20 Hz) (Chouhan et al., 2010, 2012). 30 min of stimulation induces the formation of Atg8 positive puncta. These puncta do not form when neurons are not stimulated nor in the absence of extracellular Ca^2+^ (Figure 1A-B’, E). Likewise, Atg8 labeled structures do not form when neurons are stimulated in the presence of the membrane-permeable Ca^2+^ chelator EGTA-AM (Figure 1C-E). We conclude that Ca^2+^ influx upon neuronal stimulation triggers the accumulation of Atg8-labeled autophagosomes at synaptic boutons.

**Figure 1.**
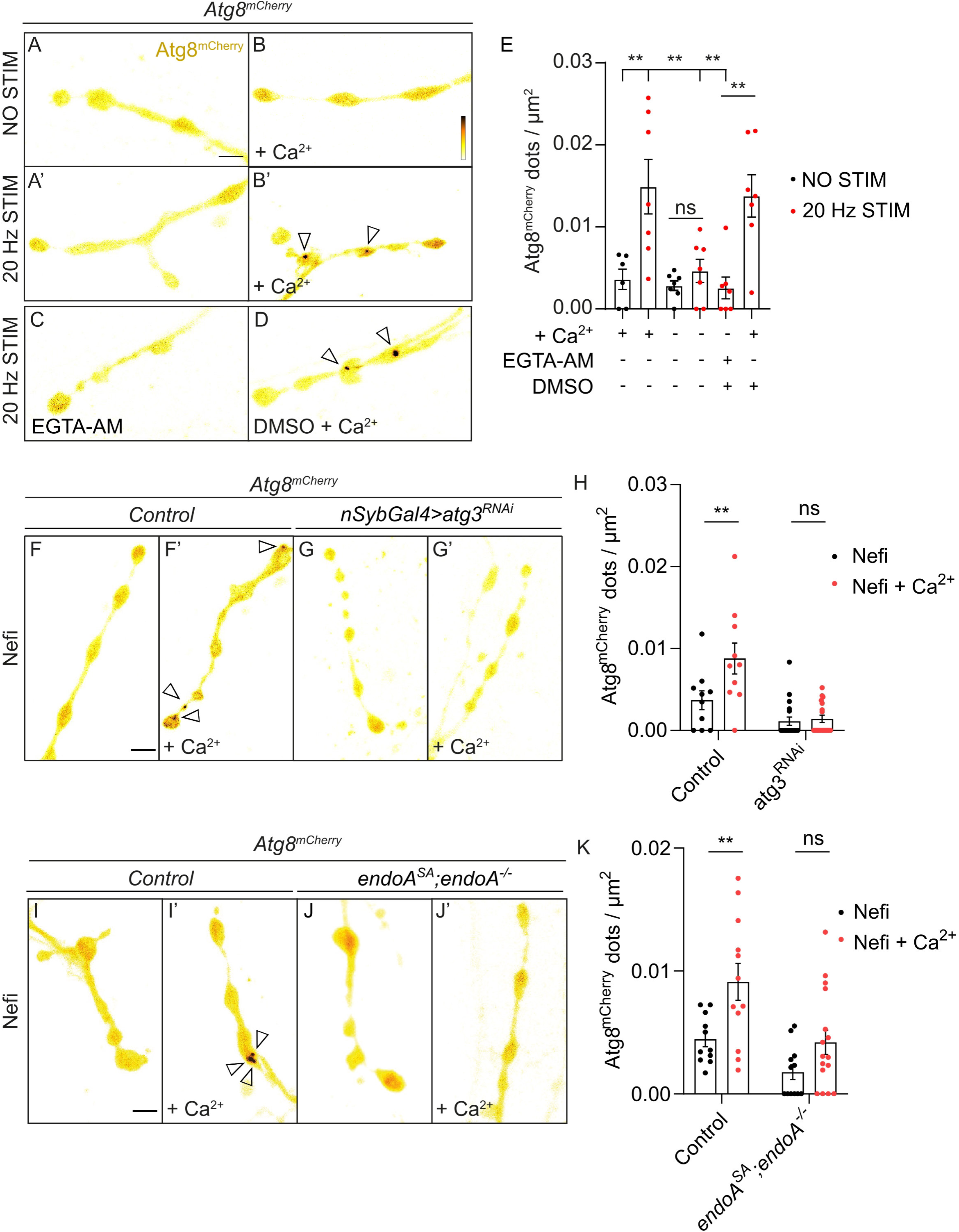
Ca^2+^ influx-induced synaptic autophagy is EndoA dependent. (A-E) Live imaging of non-stimulated and stimulated (following 30 min of 20 Hz electrical nerve stimulation) of *Drosophila* larvae NMJ boutons expressing Atg8^mCherry^ at endogenous levels in the absence of Ca^2+^ (A, A’), presence of Ca^2+^ (B, B’), presence of EGTA-AM (No Ca^2+^ in the buffer) (C) and presence of DMSO plus Ca^2+^ (D). Fluorescence intensities shown using scale (0-1292 gray value) indicated in (B). (E) Quantification of the number of Atg8^mCherry^ dots (arrowheads). Error bars represent mean ± SEM; scale bar: 5 µm. Statistical significance was calculated with an ordinary one-way ANOVA with Tukey’s multiple comparison test: \**\* P* < 0.01, ns, not significant, n ≥ 6 larvae (24 NMJs) per genotype. (F-H) Live imaging of genomically expressed Atg8^mCherry^ in NMJ boutons of *w* control animals (F, F’) and of animals expressing RNAi against *atg3* (under the control of the pan-neuronal driver *nSyb-Gal4*) (G, G’). (F, G) Non-stimulated animals were incubated for 30 min in HL3 solution containing calcium channel agonist – Nefiracetam (Nefi) (10 µM) and post-synaptic glutamate receptor blocker 1-Naphthylacetyl spermine trihydrochloride (NAS) (100 µM). (F’, G’) Stimulated animals were incubated for 30 min in HL3 solution containing Nefi (10 µM), NAS (100 µM) and CaCl_2_ (1 mM). (H) Quantification of the number of Atg8^mCherry^ dots (arrowheads). Error bars represent mean ± SEM; scale bar: 5 µm. Statistical significance was calculated with an ordinary two-way ANOVA with Šidàk multiple comparison test: \**\* P* < 0.01, ns, not significant, n ≥ 9 larvae (36 NMJs) per genotype. (I-K) Live imaging of genomically expressed Atg8^mCherry^ in NMJ boutons of *w* control animals (I, I’) and of *endoA*^-/-^ null mutant animals expressing phosphodead *endoA^S75A^* at endogenous levels (J, J’). (K) Quantification of the number of Atg8^mCherry^ dots (arrowheads). Error bars represent mean ± SEM; scale bar: 5 µm. Statistical significance was calculated with an ordinary two-way ANOVA with Šidàk multiple comparison test: \**\* P* < 0.01, ns, not significant, n ≥ 11 larvae (44 NMJs) per genotype. Full genotypes are included in the methods.

To find independent evidence for this, we used Nefiracetam (Nefi), a compound that opens L/N-type Ca^2+^ channels (Nishizaki et al., 1998; Yoshii and Watabe, 1994; Yoshii et al., 2000). First, we confirmed that Nefiracetam causes Ca^2+^ influx at *Drosophila* NMJ boutons using GCaMP6 imaging (Supplemental Figure 1A-B). We then incubated live NMJs with Nefiracetam and observed that this induces autophagy in the presence of extracellular Ca^2+^, but not when Ca^2+^ is omitted (Figures 1F, F’, H). To verify that the process induced by Nefiracetam and Ca^2+^ is indeed autophagy, we lowered the expression of the essential autophagy protein Atg3 using the expression of Atg3 RNAi in neurons (Soukup et al., 2016). Under these conditions Nefiracetam and Ca^2+^ do not cause Atg8^mCherry^ to be recruited, and the marker remains cytosolic (Figure 1G-H). Hence, Ca^2+^ influx induced by Nefiracetam induces autophagy at synapses.

EndoA is a synaptic protein involved in synaptic vesicle endocytosis (Milosevic et al., 2011; Ringstad et al., 1999; Schuske et al., 2003; Verstreken et al., 2002). Previous reports have suggested the protein is responsive to Ca^2+^, but the consequences and functional relevance of this are not known (Chen et al., 2003; Kroll et al., 2019; Yang et al., 2021; Zhang et al., 2012a). Moreover, we have previously shown that the expression of an EndoA phospho-mutant, EndoA^S75A^ instead of wild type EndoA blocks starvation-induced synaptic autophagy (Soukup et al., 2016). We therefore wondered whether EndoA would also be required for synaptic autophagy induced by Ca^2+^-influx. Interestingly, application of Nefiracetam and Ca^2+^ in EndoA^S75A^ mutant animals did not induce synaptic autophagy (Figure 1I-K). These results indicate that EndoA is required for stimulation-induced synaptic autophagy.

### The EndoA flexible region regulates Dynamin binding

Past work suggests that a negatively charged residue, E264 (in Rat ENDOA2) in the unstructured, flexible region between the BAR and SH3 domains of EndoA mediates its sensitivity to Ca^2+^ (Zhang et al., 2012a). This amino acid is evolutionary very well conserved and corresponds to D265 in *Drosophila* EndoA (Figure 2A). We mutated this residue to a neutral alanine to mimic Ca^2+^-unresponsiveness (EndoA^D265A^) and to a positively charged arginine to mimic Ca^2+^-induction (EndoA^D265R^). We expressed these mutant proteins and wild type EndoA in *E. coli*, purified them to homogeneity (Supplemental Figure 2A) and verified the integrity of our proteins by SEC-MALS (Supplemental Figure 2B). To assess if the D265 mutations cause conformational changes, as speculated in literature (Chen et al., 2003; Zhang et al., 2012a), we carried out a battery of biophysical analyses, including Fourier Transform InfraRed (FTIR) spectroscopy, Dynamic Light Scattering (DLS), Small Angle X-ray Scattering (SAXS) and assessed the proteins thermal stability (Supplemental data and Supplemental Figure 2C-H). However, none of these methodologies revealed significant structural differences between the mutant and wild type proteins. This indicates that the mutant proteins retain their ability to dimerize, do not majorly affect secondary structure composition and have similar hydrodynamic radii compared to the wild type protein. Furthermore, the D265 mutations also do not cause obvious conformational rearrangements.

**Figure 2.**
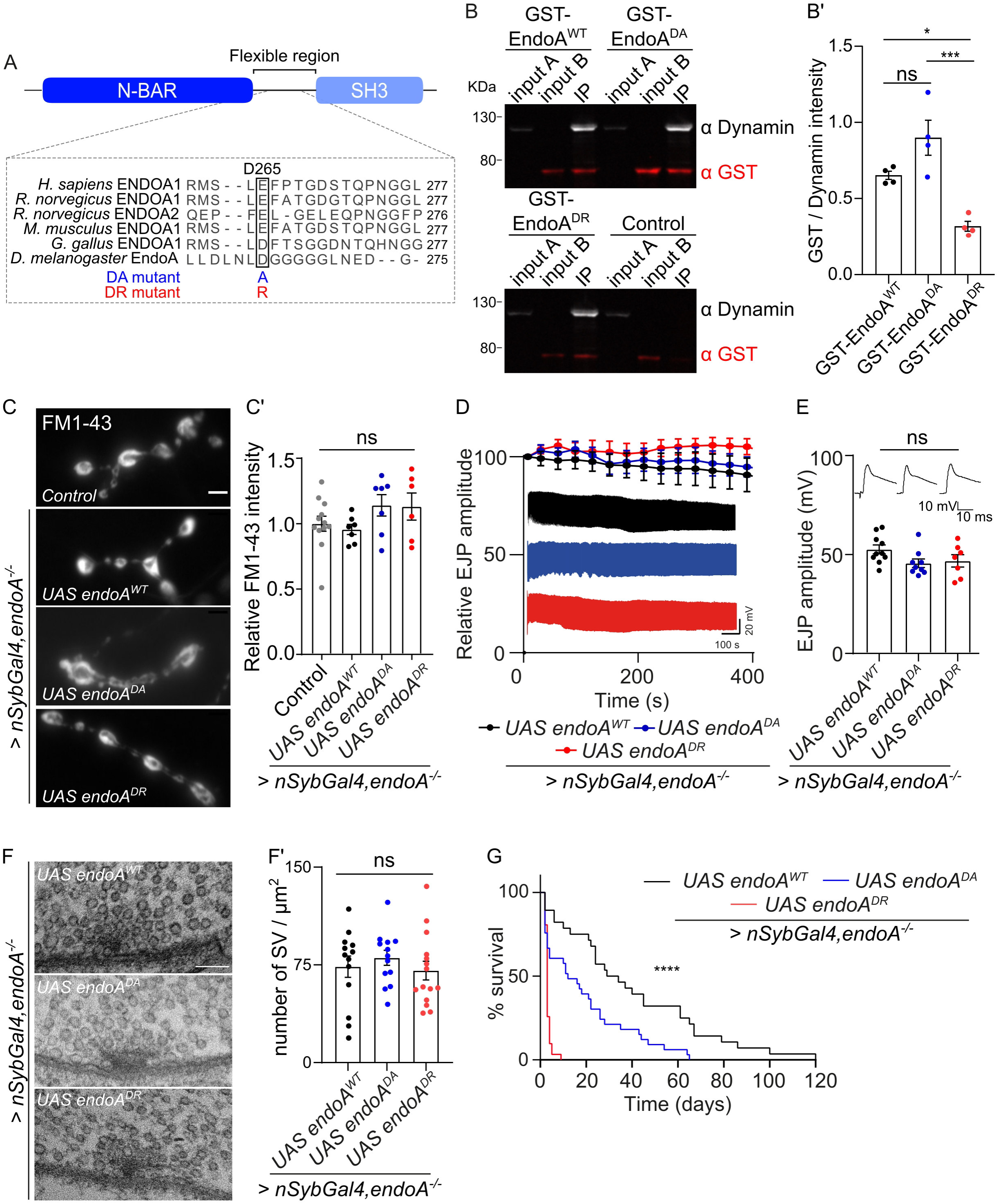
EndoA mutants increase lethality in an endocytic-independent manner. (A) Proteins alignment showing conservation of the negatively charged glutamic acid in position 264 of *Rat* ENDOA2 and negatively charged aspartic acid in position 265 of *Drosophila* EndoA flexible region. (B-B’) Co-immunoprecipitation of Dynamin and GST-EndoA wild type and mutants. (B) Western blot of 1 % control fly heads (*w^1118cs^*, input A) and 1 % purified GST-EndoA (input B) and whole co-IP fraction. Blot probed with anti-Dynamin (expected size 97 KDa) and anti-GST (expected size 67 KDa) to assess co-IP efficiency. (B’) Quantification of GST/Dynamin signal in IP lane. Error bars represent mean ± SEM. Statistical significance calculated with an ordinary one-way ANOVA with Tukey’s multiple comparison test: * *P* < 0.05, *** *P* < 0.001 ns, not significant. Experiment repeated in 4 independent replicates. (C) Representative images of boutons loaded (1 min, 90 mM KCl, 1.5 mM CaCl2) with FM 1- 43 (4 µM) and quantification (C’) of the following genotypes: control (*nSyb-Gal4/+*), *endoA*^-/-^ animals that express *endoA^WT^, endoA^D265A^* or *endoA^D265R^* under the control of *nSyb-Gal4*. Scale bar: 5 µm. Statistical significance was calculated with an ordinary one-way ANOVA with Tukey’s multiple comparison test: ns, not significant, n ≥ 6 larvae (24 NMJs) per genotype. (D) Relative EJP amplitudes and raw traces recorded during 400 s of 10 Hz stimulation train of the indicated genotypes. (E) Quantification of EJP amplitudes of indicated genotypes. Statistical significance was calculated with an ordinary one-way ANOVA with Tukey’s multiple comparison test: ns, not significant, n ≥ 7 larvae per genotype. (F) Representative electron micrographs of NMJ bouton and (F’) quantification of the number of synaptic vesicles (SV) per bouton area (/ μm^2^) for animals of the indicated genotypes. Scale bar: 150 nm. Data point represent single boutons from ≥ 3 animals per genotype. Statistical significance calculated with an ordinary one-way ANOVA with Tukey’s multiple comparison test: ns, not significant. (G) Survival rate (in percentage) of adult *endoA*^-/-^ *Drosophila* expressing *endoA^WT^, endoA^D265A^*, or *endoA^D265R^* under the control of the pan-neuronal driver *nSyb-Gal4* over 120 days. Median survival of *endoA^WT^* expressing flies is 31.5 days, 12 days for *endoA^D265A^* expressing flies and 3 days for *endoA^D265R^* expressing flies. Statistical significance was calculated using Log-rank Mantel-Cox test: \*\*\**\* P* < 0.0001, n ≥ 28 animals per genotype.

The E264 in the flexible region of rat ENDOA2 is thought to affect the binding efficiency of proteins to the EndoA-SH3 domain. We therefore assessed the ability of EndoA^D265R^ and EndoA^D265A^ to bind Dynamin, a well-known EndoA-interaction partner that mediates synaptic vesicle endocytosis at the plasma membrane (Ringstad et al., 1999). We found that *Drosophila* EndoA^D265R^ binds significantly less Dynamin than EndoA^WT^, and that EndoA^D265A^ binds more Dynamin than EndoA^D265R^ (Figure 2B-B’). These findings indicate that D265 regulates the association of EndoA with Dynamin, one of its major binding partners.

### D265 in EndoA mediates Ca^2+^-induced synaptic autophagy

To test if D265 mediates the Ca^2+^ responsiveness of EndoA in autophagy, we generated transgenic flies and expressed EndoA^D265A^, EndoA^D265R^ or wild type EndoA using a pan-neuronal driver (*nSyb-Gal4*) in *endoA^-/-^* null mutants. We show that our conditions result in very similar expression levels to endogenous EndoA expression in control animals (Supplemental Figure 3A, A’) and that the proteins localize to presynaptic terminals (see below). Next, we assessed if these mutant EndoA proteins can recapitulate *in vivo* functions of the wild type protein. Unlike *endoA^-/-^* mutants that die as pupae, neuronal expression of EndoA^D265A^ or EndoA^D265R^ rescues lethality and adult flies emerge (Supplemental Figure 3B). However, most fail to expand their wings, suggesting the animals are weak (Supplemental Figure 3C-C’). Additionally, *endoA^-/-^* animals that express EndoA^D265A^ or EndoA^D265R^ are extremely uncoordinated and the animals die much earlier than *endoA^-/-^* mutants that express wild type EndoA (Figure 2G). Hence, these data suggest that mutations within EndoA flexible region fundamentally affect EndoA function.

*EndoA^-/-^* null mutants show a severe defect in synaptic vesicle endocytosis (Dickman et al., 2005; Guichet et al., 2002; Verstreken et al., 2002). To assess whether EndoA^D265A^ and EndoA^D265R^ affect synaptic vesicle cycling, we performed a FM 1-43 dye uptake assay. Larval fillets were incubated with the lipophilic dye FM 1-43, which is internalized into newly formed synaptic vesicles upon nerve stimulation (Ramaswami et al., 1994). *EndoA^-/-^* animals expressing EndoA^D265A^ or EndoA^D265R^ show efficient FM 1-43 dye uptake that is similar to that measured in *endoA^-/-^* mutants that express wild type EndoA (Figure 2C-C’). Endocytic defects in *endoA^-/-^* mutants also cause a decline in neurotransmitter release during prolonged 10 Hz stimulation (Verstreken et al., 2002, 2003). To test if this response was affected in D265 mutant flies, excitatory junctional potentials (EJPs) were recorded. The EJP amplitude during low frequency stimulation (0.2 Hz-1 Hz) and during high frequency stimulation (10 Hz, 400 s) was very similar across our conditions (Figure 2D-E). Moreover, defects in endocytosis in *endoA^-/-^* mutants severely affect vesicle replenishment at synaptic terminals (Guichet et al., 2002; Verstreken et al., 2002, 2003). Therefore, we conducted transmission electron microscopy (TEM) to reveal the ultrastructure of NMJ boutons, but did not find significant differences in synaptic vesicle number per area (Figure 2F-F’). We conclude that the D265 EndoA mutants do not affect synaptic vesicle endocytosis in a significant manner, but cause adult flies to be uncoordinated resulting in early death (Figure 2G). This suggests that important EndoA functions, other than endocytosis, are affected by D265.

In light of these results, we asked if Ca^2+^ influx-induced synaptic autophagy might be affected in EndoA^D265A^ and EndoA^D265R^ mutants. We determined the distribution of Atg8^mCherry^ in the mutant animals in response to Ca^2+^ influx. In non-induced conditions (absence of extracellular Ca^2+^), Atg8^mCherry^ remains cytosolic and does not accumulate in puncta in *endoA^-/-^* mutants that express EndoA^WT^ or EndoA^D265A^ (Figure 3A-C, E). In contrast, Atg8^mCherry^-labeled puncta do form at uninduced NMJs of *endoA^-/-^* mutants that express EndoA^D265R^ (Figure 3D-E). The autophagy levels in uninduced EndoA^D265R^ mutants are similar to those observed in controls treated with Nefiracetam and Ca^2+^ (induced), indicating that autophagy is constitutively induced in EndoA^D265R^ mutants (Figure 3E). Conversely, Atg8^mCherry^ puncta fail to be formed at induced NMJs of *endoA^-/-^* mutants that express EndoA^D265A^, indicating that in this mutant autophagy is blocked (Figure 3A’-D’, E).

**Figure 3.**
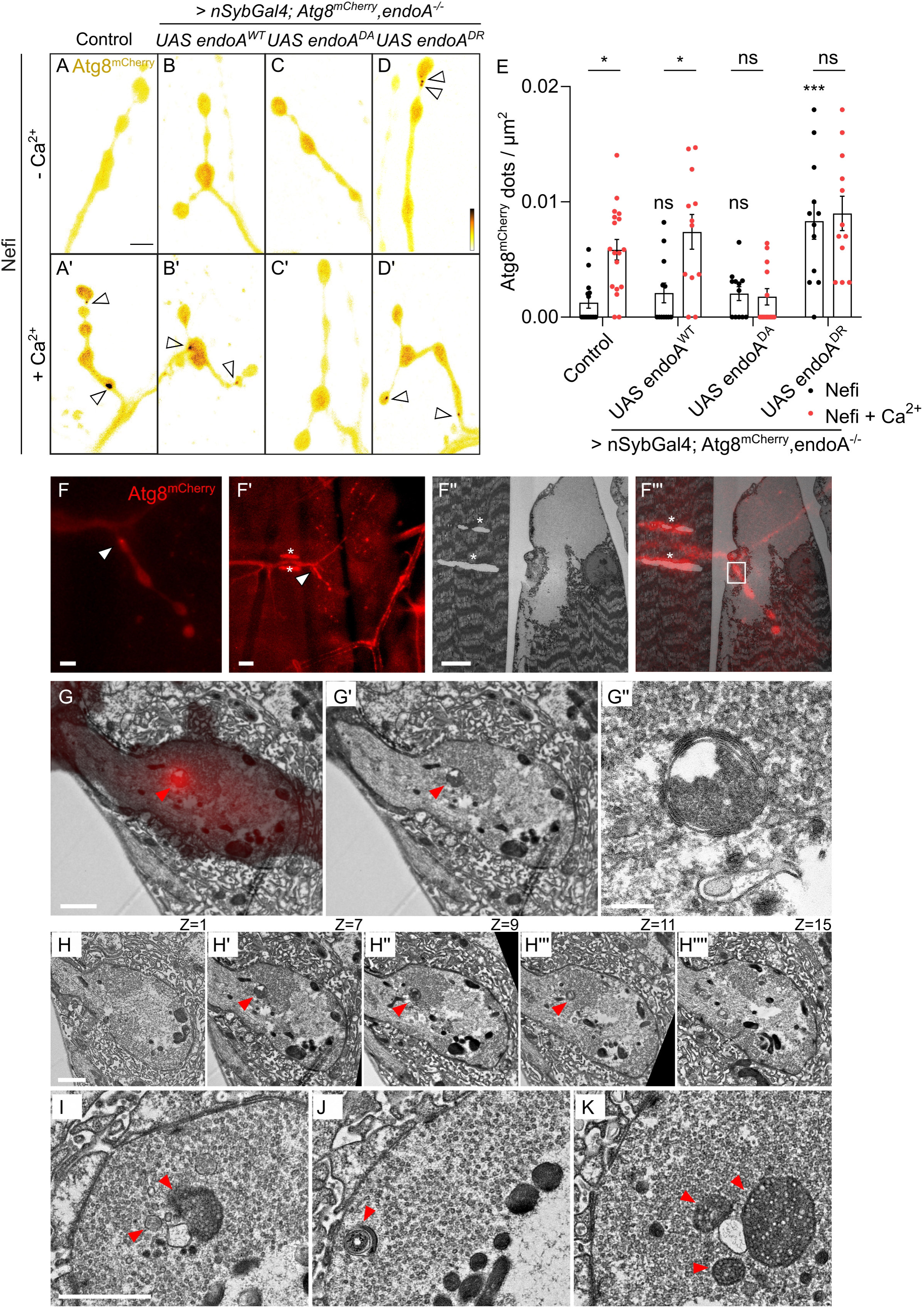
EndoA mutants alter Ca^2+^ influx mediated synaptic autophagy induction. (A-E) Live imaging of genomically expressed Atg8^mCherry^ in NMJ boutons of control (*nSyb- Gal4/+*) animals (A, A’) and of *endoA*^-/-^ animals expressing *endoA^WT^* (B, B’), *endoA^D265A^* (C, C’), and *endoA^D265R^* (D, D’) (under the control of the pan-neuronal driver *nSyb-Gal4*). Non- stimulated animals were incubated for 30 min in HL3, Nefi (10 µM) and NAS (100 µM) solution (A, B, C, D). Stimulated animals were incubated for 30 min in HL3 solution containing Nefi (10 µM), NAS (100 µM) and CaCl_2_ (1 mM) (A’, B’, C’, D’). Fluorescence intensities shown using scale (0-23645 gray value) indicated in (D). Arrowheads indicate Atg8^mCherry^ accumulations. Scale bar: 5 μm. (E) Quantification of the number of Atg8^mCherry^ dots (arrowheads) per NMJ area. Error bars represent mean ± SEM. Statistical significance was calculated with an ordinary two-way ANOVA with Tukey’s multiple comparison test: *\* P* < 0.05, \*\**\* P* < 0.001, ns, not significant, n ≥ 12 larvae (48 NMJs) per genotype. Significance levels displayed above ‘Nefi’ columns refer to comparison with ‘Nefi’ treatment (unstimulated) on control animals. (F-K) CLEM of boutons of *endoA*^-/-^ animals expressing *endoA^D265R^* under the pan-neuronal driver *nSyb-Gal4*, as well as Atg8^mCherry^ expressed at endogenous levels. (F) Single confocal slice of an example NMJ displaying an Atg8^mCherry^ structure (arrowhead). Scale bar: 5 μm. (F’) Zoom out of the same NMJ shown in (F). Asterisks indicate branding marks, arrowhead indicates Atg8^mCherry^ structure. Scale bar: 20 μm. (F’’) Electron micrograph of the same region as in (F’). Asterisks indicate branding marks, arrowhead indicates Atg8^mCherry^ structure. Scale bar: 10μm. (F’’’) Overlay of confocal image in (F’) with electron micrograph in (F’’). Asterisks indicate branding marks, arrowhead indicates Atg8^mCherry^ structure. Scale bar: 10 μm. (G-G’) Zoomed representation of the bouton containing the Atg8^mCherry^ structure shown as overlay (G) and electron micrograph only (G’). Red arrowhead indicates the structure correlating with the mCherry signal. Scale bar: 1 μm. (G’’) Magnification of the putative autophagosome. Scale bar: 200 nm. (H-H’’’’) Single TEM slices showing the putative autophagosomal structure visible in multiple consecutive slices (red arrowhead). Z=1: 0, Z=7: 42 nm, Z=9: 560 nm, Z=11: 700 nm, Z=15: 980 nm. Scale bar: 1 μm. (I-K) Examples of autophagosomal structures (red arrowheads) from the same animal. Scale bar: 1 μm.

To reveal the ultrastructure of the Atg8^mCherry^-labeled structures, we resorted to correlative light and electron microscopy (CLEM). First, we imaged by confocal microscopy Atg8^mCherry^ puncta in NMJs of *endoA^-/-^* animals that express EndoA^D265R^ and used a two-photon laser to create branding marks around the region of interest. We used these to identify relevant boutons in block face scanning EM and processed the samples for imaging by TEM. We then manually overlayed confocal light microscopy sections with the TEM images (Soukup et al., 2016). We found that the Atg8^mCherry^ puncta overlap with structures reminescent of degradative organelles including lysosomes and autophagosomes (Figure 3F-K). This confirms the autophagosome/degradative nature of the Atg8^mCherry^ labeled organelles that form in EndoA^D265R^ animals. Taken together, these results indicate that EndoA D265 mutants are unresponsive to Ca^2+^-influx and that EndoA^D265R^ consitutively induces autophagy at synapses, while EndoA^D265A^ is inert and fails to induce the process even upon Ca^2+^ influx.

### Ca^2+^ influx affects the nanoscale organization of EndoA at synapses

To understand how these EndoA mutants differentially affect autophagy at synapses, we determined the localization of wild type and mutant EndoA proteins at live synapses with and without induction by Nefiracetam and Ca^2+^. We took advantage of the nanoscale resolution provided by the Airy scan detector in confocal laser-scanning microscopy (Huff, 2015). Ca^2+^ influx causes a redistribution of endogenous EndoA or EndoA^WT^ neuronally expressed in *endoA^-/-^* mutants (single confocal slices in Figure 4A-F). Wild type EndoA moves from its preferential peripheral localization towards the synaptic bouton lumen (Figures 4A–A’, C-C’ G). Interestingly, EndoA^D265A^ remains mostly at the bouton periphery even upon Ca^2+^ influx (Figure 4B-B’, E, G). Conversely, EndoA^D265R^ is distributed across the bouton already at rest (Figure 4D-G). In contrast to wild type EndoA, the distribution of EndoA^D265A^ or EndoA^D265R^ does not change upon Ca^2+^-influx, confirming these mutants are Ca^2+^ insensitive. We next asked whether the redistribution of EndoA would cause increased localization of the protein close to Atg8^mCherry^ labeled autophagosomes. We therefore expressed Atg8^mCherry^ and quantified the amount of EndoA in the 100 nm zone around specified Atg8^mCherry^ puncta. As predicted, EndoA localizes more in the proximity of Atg8^mCherry^ labeled structures in response to Ca^2+^ influx and when harboring the D265R mutation. Conversely, significantly less EndoA localizes around (rare) Atg8^mCherry^ labeled structures in unstimulated boutons, or upon expression of EndoA^D265A^ (Figure 4H-H’). These data suggest that increased Ca^2+^ influx enables EndoA re-distribution, including to synaptic autophagosomes.

**Figure 4.**
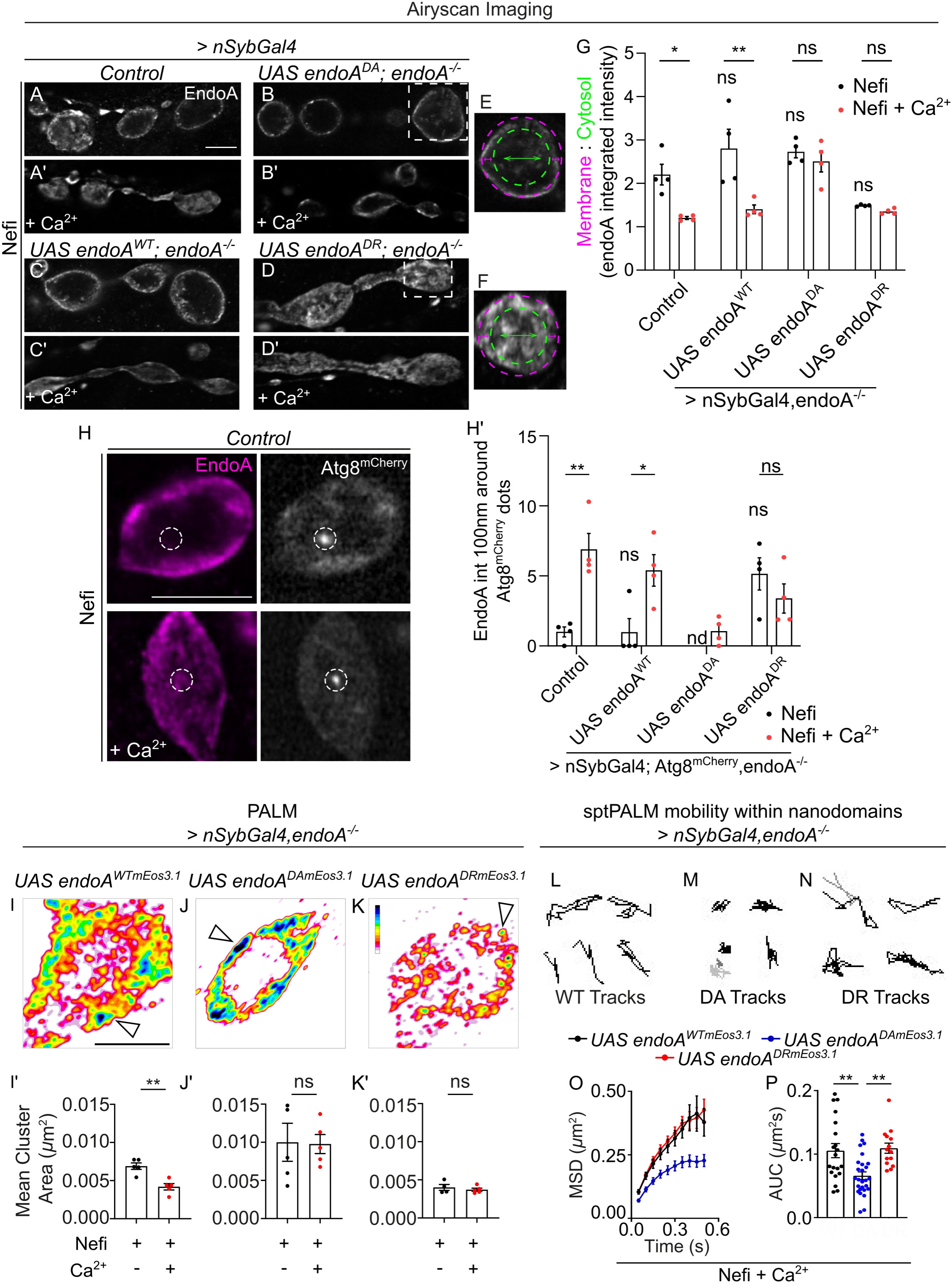
Synaptic nanoscale organization of EndoA is Ca^2+^ influx dependent. (A-G) Representative images of Airyscan confocal single slice sections of synaptic boutons of paraformaldehyde-fixed control (*nSyb-Gal4/+*) (A, A’) and *endoA*^-/-^ larvae expressing *endoA^D265A^* (B, B’)*, endoA^WT^* (C, C’), and *endoA^D265R^* (D, D’) (under the control of the pan-neuronal driver *nSyb-Gal4*) labelled with anti-EndoA antibody. Scale bar: 5µm. Non-stimulated animals were incubated for 30 min in HL3, Nefi (10 µM) and NAS (100 µM) solution (A, B, C, D). Stimulated animals were incubated for 30 min in HL3 solution containing Nefi (10 µM), NAS (100 µM) and CaCl_2_ (1 mM) (A’, B’, C’, D’). (E, F) Zoomed in synaptic boutons (from B and D respectively) showing quantification of EndoA intensity around the synaptic plasma membrane (magenta) and within the synaptic lumen - cytosol (green). See Methods for details. (G) Quantification of the EndoA integrated intensity across genotypes indicated in (A-D) showing ratio of EndoA intensity at the membrane to that within the cytosol. Error bars represent mean ± SEM; statistical significance was calculated with an ordinary two-way ANOVA with Tukey’s multiple comparison test: *\* P* < 0.05, \**\* P* < 0.01, ns, not significant, n ≥ 4 larvae (16 NMJs) per genotype. Significance levels displayed above ‘Nefi’ columns refer to comparison with ‘Nefi’ treatment (unstimulated) on control animals. (H-H’) Representative images of Airyscan confocal single slice sections of individual synaptic boutons of paraformaldehyde-fixed control (*nSyb-Gal4/+*) in stimulated and non stimulated conditions, labelled with anti-EndoA antibody and expressing Atg8^mCherry^ at endogenous levels. Scale bar: 5 µm. Only synaptic boutons with Atg8^mCherry^ punctae (enclosed in dotted circles) indicative of autophagosomes were selected and EndoA integrated density in a radius of 100 nm from the Atg8^mCherry^ punctae was measured. (H’) Quantification of the normalized integrated EndoA intensity across genotypes indicated in (A-D) showing EndoA intensity 100 nm around Atg8^mCherry^ punctae. Data from all genotypes was normalized to control (*nSyb- Gal4/+*) unstimulated data. Error bars represent mean ± SEM; statistical significance was calculated with an ordinary two-way ANOVA with Tukey’s multiple comparison test: *\* P* < 0.05, \**\* P* < 0.01, ns, not significant, nd, no data, n ≥ 4 larvae (16 NMJs) per genotype. (I-K) Transgenic *endoA*^-/-^ larvae expressing *endoA^WT^*^::mEos3.1^, *endoA^D265A^*^::mEos3.1^ or *endoA^D265R^*^::mEos3.1^ (under the control of the pan-neuronal driver *nSyb-Gal4*) were imaged using single molecule localization photoactivated localization microscopy (PALM) at 20 Hz. Representative images show cluster map colour-coded for cluster size and density distribution of *endoA*^::mEos3.1^ generated by density-based spatial clustering of applications with noise (DBSCAN) analysis. Arrowheads indicate EndoA nanodomains. Fluorescence intensity shown using indicated scale (2-203). Scale bar: 2 µm. (I’-K’) Quantification of the mean cluster area of *endoA^WT^*^::mEos3.1^, *endoA^D265A^*^::mEos3.1^ and *endoA^D265R^*^::mEos3.1^ in non-stimulated and stimulated conditions. Error bars represent mean ± SEM; statistical significance was calculated with an student *t-*test two-tailed unpaired distribution: \**\* P* < 0.01, ns, not significant, n ≥ 5 larvae (20 NMJs) per genotype. (L-N) Transgenic *endoA*^-/-^ larvae expressing *endoA^WT^*^::mEos3.1^, *endoA^D265A^*^::mEos3.1^ or *endoA^D265R^*^::mEos3.1^ (under the control of the pan-neuronal driver *nSyb-Gal4*) were imaged using single particle tracking photoactivated localization microscopy (sptPALM) at 20 Hz. Representative trajectories located within EndoA nanodomains in NMJ boutons of *endoA*^-/-^ larvae expressing *endoA^WT^*^::mEos3.1^ (L), *endoA^D265A^*^::mEos3.1^ (M) and *endoA^D265R^*^::mEos3.1^ (N). This is quantified for each genotype as mean square displacement (MSD) as a function of time (O). (P) Quantification of the area under the MSD curve (µm^2^s) represented in (O). Error bars represent mean ± SEM; statistical significance was calculated with an ordinary one-way ANOVA with Tukey’s multiple comparison test: \**\* P* < 0.01, n ≥ 4 larvae (n ≥ 14 nanodomains) per genotype.

To further understand the movements of EndoA at the bouton periphery we resorted to photoactivated localization microscopy (PALM) under oblique illumination that was developed for *Drosophila* NMJs (Bademosi et al., 2017, 2018; Vanhauwaert et al., 2017). This technology allows the tracking of proteins tagged with photoconvertible fluorescent probes at <50 nm resolution and within 200-400 nm proximity to the synaptic plasma membrane (Tokunaga et al., 2008). We tagged wild type, EndoA^D265A^ and EndoA^D265R^ mutants with mEOS3.1 and expressed the proteins in *endoA^-/-^* mutants. mEOS is a photoconvertible fluorescent protein whose stochastic change in emission spectrum from green to red allows for single molecule localization (Manley et al., 2008; Zhang et al., 2012b). Animals were or were not stimulated with Nefiracetam and Ca^2+^, fixed and analyzed by PALM (Supplementary Figure 4A-A’’’). Interestingly, wild type EndoA^WT::mEos3.1^ organizes in ‘hot-spots’ or ‘nanometer-sized cluster of proteins – nanodomains’ at the bouton periphery (Supplementary Figure 4A’’’). These EndoA^WT::mEos3.1^ nanodomains are significantly smaller when NMJs are stimulated or when imaging EndoA^D265R::mEos3.1^ as compared to unstimulated samples or when imaging EndoA^D265A::mEos3.1^ (Figure 4I’-K’, Supplemental Figure 4B-D’). Our data are consistent with the idea that EndoA^D265A^ interacts with other proteins in the periphery including Dynamin (Figure 2B-B’) and is thus more confined within larger nanodomains located juxtamembrane, while EndoA^D265R^ is more localized to the synaptic lumen and thus away from the nanodomains (Figure 4D-D’,G).

Finally, we characterized the re-arrangements of EndoA in nanodomains upon Ca^2+^ influx using live single particle tracking PALM (sptPALM) (Supplemental Figure 5A-B’) (Manley et al., 2008). We incubated NMJs with Nefiracetam and Ca^2+^ and tracked individual EndoA^WT::mEos3.1^, EndoA^D265A::mEos3.1^ or EndoA^D265R::mEos3.1^ molecules (neuronally expressed in *endoA^-/-^*) and plotted their trajectory map (Figure 4L-N; Supplemental Video 1). We first confirmed the existence of EndoA-nanodomains at the bouton periphery using the high-resolved intensity maps that are generated during the sptPALM processing (Supplemental Figure 5C–C’). Then, we assessed the mobility behavior of EndoA in these nanodomains by analyzing the mean square displacement (MSD) of the single proteins within the nanodomains. This parameter reveals the level of confinement of motion. Upon Ca^2+^ influx, EndoA^D265A::mEos3.1^ is significantly more confined within these nanodomains than EndoA^WT::mEos3.1^ or EndoA^D265R::mEos3.1^ (Figure 4O-P). We reach a similar conclusion when analyzing the instantaneous diffusion coefficients of EndoA across entire NMJs, revealing increased mobility of EndoA^WT::mEos3.1^ in response to Ca^2+^ influx, while EndoA^D265R::mEos3.1^ and EndoA^D265A::mEos3.1^ are insensitive (Supplemental Figure 5D-F’’’). Mechanistically, these results indicate that the D265A mutation confines EndoA in nanodomains at the bouton periphery and that Ca^2+^-influx alters the diffusion of the wild type protein to localize to the bouton lumen.

### EndoA D265 mutations cause neurodegeneration

Having characterized the involvement of EndoA in synaptic autophagy, we turned to examine its potential implication in neurodegeneration as defects in autophagy are frequently connected to these disorders. We specifically tested if defects in stimulation-induced synaptic autophagy lead to neuronal demise by assessing neuronal integrity in *endoA^-/-^* mutants expressing EndoA^D265R^, EndoA^D265A^ or EndoA^WT^. We started by placing animals for 3 or 7 days in constant dark to avoid retinal stimulation and then used these animals to record electroretinograms (ERGs). ERGs measure the depolarization of photoreceptors in response to a brief (1 s) light pulse. When photoreceptor neurons degenerate, the amplitude of this depolarization is reduced (Chouhan et al., 2016; Soukup et al., 2016; Wang and Montell, 2007). While the amplitude of depolarization in 3-day-old and 7-day-old control and EndoA^D265A^ mutant flies is similar, EndoA^D265R^ already show a clear age-dependent reduction in ERG amplitude when kept in dark conditions (Figure 5A, A’, C, C’, E, E’, G, G’, I). We also placed the flies in constant light to cause light-induced stress and repeated this experiment. While control and *endoA^-/-^* flies expressing EndoA^WT^ are fine, now both *endoA^-/-^* flies expressing EndoA^D265A^ and EndoA^D265R^ show strongly reduced ERG depolarization amplitudes (Figure 5A’’,C’’,E’’,G’’,I), a hallmark of neurodegeneration. To further corroborate these findings we also evaluated neuronal morphology in toluidine blue-stained retinal sections. Quantification of the number of intact ommatidia with 7 visible photoreceptors validated the ERG data and also shows age-dependent degeneration in 7-day old dark-reared EndoA^D265R^-expressing flies and in EndoA^D265R^- and EndoA^D265A^-expressing flies kept for 7 days in constant light (Figure 5B-H’’, J). These data further indicate that expression of EndoA^D265R^ is a more severe condition than EndoA^D265A^, but that both EndoA^D265R^ and EndoA^D265A^ cause neurodegeneration. They also provide evidence that a tight balance of stimulation-induced autophagy under control of EndoA is critical for neuronal survival.

**Figure 5.**
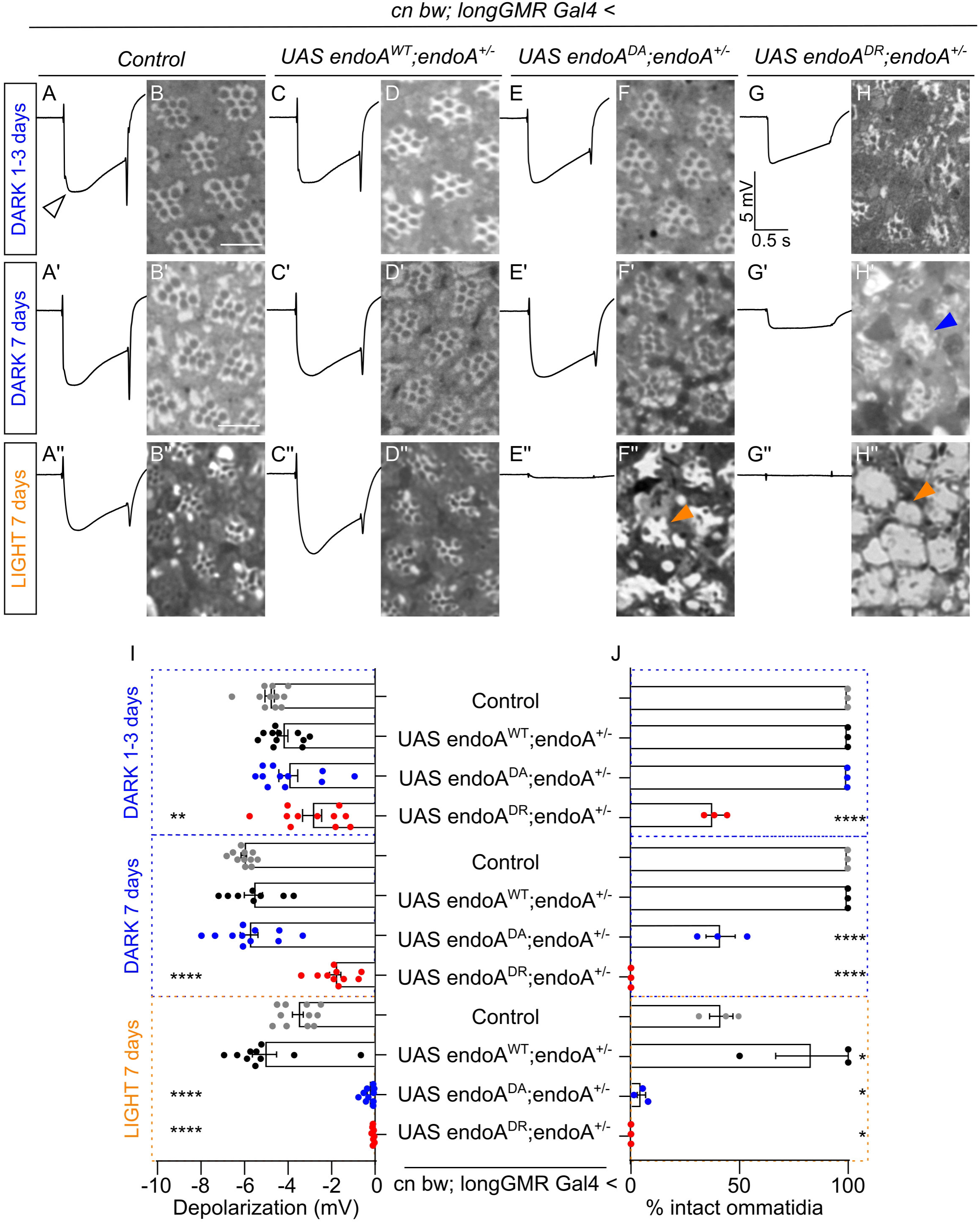
EndoA insensitivity to Ca^2+^ influx induces neurodegeneration. (A-G’’) Representative ERG traces recorded from control (*cn bw; longGMR Gal4/+*) and *endoA*^+/-^ mutant flies expressing *endoA^WT^*, *endoA^D265A^* or *endoA^D265R^* under a photoreceptor specific driver (*longGMR-Gal4*). *Cn bw* mutations remove the protective eye pigmentation. Prior to ERG recording, animals were exposed to 1-3 (A, C, E, G) or 7 days of constant dark (A’, C’, E’, G’) or 7 days of constant light (A’’, C’’, E’’, G’’). Average traces are depicted in black. Arrowhead indicates ERG depolarization. (I) Quantification of ERG depolarization amplitude recorded upon a 1 s light pulse in flies exposed to dark or light. Plotted bars show mean ± SEM. Statistical significance calculated with an ordinary one-way ANOVA with Tukey’s multiple comparison test: *\* P* < 0.05, *\*\* P* < 0.01, \*\*\**\* P* < 0.0001. Plotted significance level refers to the difference to the control (*cn bw; longGMR Gal4/+*) of the indicated condition (light or dark). Number of recorded flies per condition ≥ 8. (B-H’’) Histological sections of retinas of flies exposed for 1-3 (B, D, F, H) or 7 days to constant dark (B’, D’, F’, H’) or constant light (B’’, D’’, F’’, H’’) stained with toluidine blue. Arrowheads indicate morphologically abnormal ommatidia. Scale bar: 10 µm. (J) Quantification of the number of intact ommatidia (expressed in % of the total) meant as ommatidia in which all 7 rhabdomeres are visible. Plotted bars show mean ± SEM. Statistical significance calculated with an ordinary one-way ANOVA with Tukey’s multiple comparison test: *\* P* < 0.05, *\*\* P* < 0.01, \*\*\**\* P* < 0.0001. Plotted significance level refers to the difference to the control (*cn bw; longGMR Gal4/+*) of the indicated condition (light or dark). Single data points represent the average % of intact ommatidia of 3 histological sections of the same animal. Analyzed animals: 3.

### A Parkinson’s disease risk variant in *SH3GL2* impairs Ca^2+^-induced synaptic autophagy

There are several lines of human genetic evidence that indicate a role for ENDOA1 in the development of PD. There are two independent GWAS signals in the vicinity of the *SH3GL2* gene that encodes ENDOA1 (Alfradique-Dunham et al., 2021; Chang et al., 2017; Germer et al., 2019; Nalls et al., 2019). Furthermore, there is a risk-conferring missense mutation in the *SH3GL2* gene of PD patients (Germer et al., 2019) that encodes a Glycine to Valine transition at position 276 in the flexible region of ENDOA1. Given the proximity of G276V to the autophagy-controlling D265 position (E264 in ENDOA1) (Figure 6A), we speculated that this risk mutation might also affect Ca^2+^ influx-induced synaptic autophagy.

**Figure 6.**
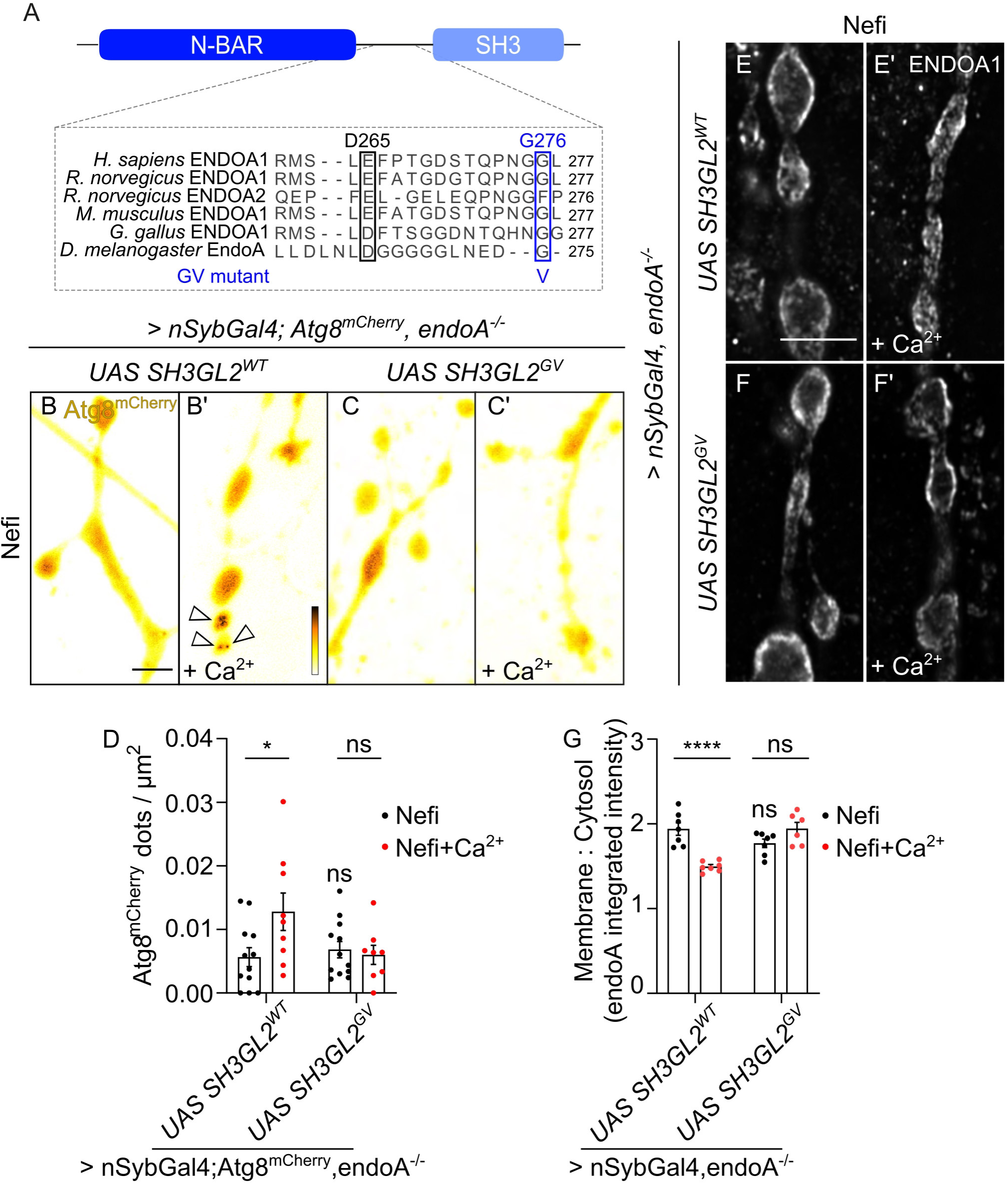
*SH3GL2* Parkinson’s disease coding variant impairs Ca^2+^-induced synaptic autophagy. (A) Protein alignment showing evolutional conservation of glycine 276 on the flexible region of *Human* ENDOA1 and the proximity to the aspartic acid at position 265 (corresponding to glutamic acid 264 in *Human* ENDOA1). (B-C’) Representative live confocal images of NMJ boutons of *endoA^-/-^* animals expressing Atg8^mCherry^ at endogenous level and *SH3GL2^WT^* or *SH3GL2^G276V^* under a pan-neuronal driver (*nSyb-Gal4*). Non-stimulated animals were incubated for 30 min in HL3, Nefi (10 µM) and NAS (100 µM) solution (B, C). Stimulated animals were incubated for 30 min in HL3 solution containing Nefi (10 µM), NAS (100 µM) and CaCl_2_ (1 mM) (B’, C’). Fluorescence intensities shown using scale (0-65535 gray value) indicated in (B’). Arrowheads indicate Atg8^mCherry^ positive autophagosomes. Scale bar: 5 μm. (D) Quantification of the number of Atg8^mCherry^ dots per NMJ area. Error bars represent mean ± SEM. Statistical significance was calculated with an ordinary two-way ANOVA with Tukey’s multiple comparison test: *\* P* < 0.05, ns, not significant, n ≥ 8 larvae (32 NMJs) per genotype. Significance level displayed above ‘Nefi’ column refers to comparison with ‘Nefi’ treatment (unstimulated) on control animals. (E-F’) Representative images of Airyscan confocal single slice sections of synaptic boutons of *endoA*^-/-^ larvae expressing *SH3GL2^WT^* (E, E’) and *SH3GL2^G276V^* (F, F’) (under the control of the pan-neuronal driver *nSyb-Gal4*) labelled with anti-ENDOA1 antibody. Scale bar: 5 µm. (G) Quantification of the ENDOA1 integrated intensity across genotypes showing ratio of ENDOA1 intensity at the membrane to that within the cytosol. Error bars represent mean ± SEM; statistical significance was calculated with an ordinary two-way ANOVA with Tukey’s multiple comparison test: *\*\**\**\* P* < 0.0001, ns, not significant, n ≥ 6 larvae (24 NMJs) per genotype. Significance levels displayed above ‘Nefi’ column refer to comparison with ‘Nefi’ treatment (unstimulated) on control animals.

We first verified that the G276V mutation in human ENDOA1 does not destabilize the protein. We produced recombinant ENDOA1^WT^ and ENDOA1^G276V^ and analyzed them by SAXS. This indicates that there are no large conformational differences between them (Supplemental Figure 6A) and that the proteins are both well-folded (Supplemental Figure 6B). Next, we generated *endoA^-/-^* flies expressing human ENDOA1^WT^ and ENDOA1^G276V^ pan-neuronally. Western blot analysis confirms the proteins are expressed at similar levels (Supplemental Figure 6C-C’) and they are localized to synapses (see below). We also assessed synaptic vesicle endocytosis by measuring the uptake of FM 1-43 during stimulation. Consistent with the other tested mutations in the flexible region, we found no significant difference (Supplemental Figure 6D-D’). Together, these results indicate that the G276V mutation does not destabilize ENDOA1, that human ENDOA1 can functionally replace *Drosophila* EndoA when expressed in flies and that the G276V mutation does not affect synaptic vesicle endocytosis under the tested conditions.

To test if Ca^2+^ influx-induced autophagy is affected by the G276V pathogenic mutation, we quantified the number of Atg8^mCherry^ puncta upon induction by Ca^2+^ influx. While *endoA^-/-^* mutants expressing wild type human ENDOA1^WT^ show a significant increase in Atg8^mCherry^- labeled autophagosomes upon induction (Figure 6B-B’, D), ENDOA1^G276V^ expressing animals fail to upregulate autophagy even when induced (Figure 6C-D). We next also tested the localization of the human protein at *Drosophila* synapses using Airy scan confocal imaging. While Ca^2+^-influx causes ENDOA1^WT^ to redistribute from the periphery to a more lumenal localization (Figure 6E-E’, G), ENDOA1^G276V^ remaines confined to the periphery (Figure 6F-G). Collectively these data indicate that the G276V mutation in ENDOA1 impairs Ca^2+^ influx- induced synaptic autophagy.

To assess the validity of our result in human cells, we investigated the role of the ENDOA1^G276V^ mutation in induced human dopaminergic neurons. Therefore, we generated ENDOA1^G276V^ homozygous knock-in mutations by CRISPR/Cas9-facilitated homologous recombination in Ctrl65 “wild type” (Baumann et al., 2021) induced pluripotent stem cells (iPSCs) (Figure 7A-A’, Supplemental Figure 7A-B). Then, ventral midbrain dopaminergic neurons (vmDAn) were produced using an optimized protocol for midbrain floor plate differentiation (Supplemental Figure 7C). Terminal differentiation *in vitro* until day 55-60 ultimately yields ∼50% of TH- positive neurons for both unedited and ENDOA1^G276V^ mutant cells (Figure 7B-B’). Finally, we assessed autophagy by quantifying the number of LC3B puncta (by antibody labeling) within TH-labeled neurites. This antibody labels lipidated LC3, the human orthologue of Atg8a (Klionsky et al., 2021; Martinet et al., 2013). We observed significantly less autophagosomes in ENDOA1^G276V^ mutant vmDAn compared to control neurons (Figure 7C-C’). These results support the conclusion that the PD risk-conferring variant ENDOA1^G276V^ impaires autophagy in neurites of induced vmDAn.

**Figure 7.**
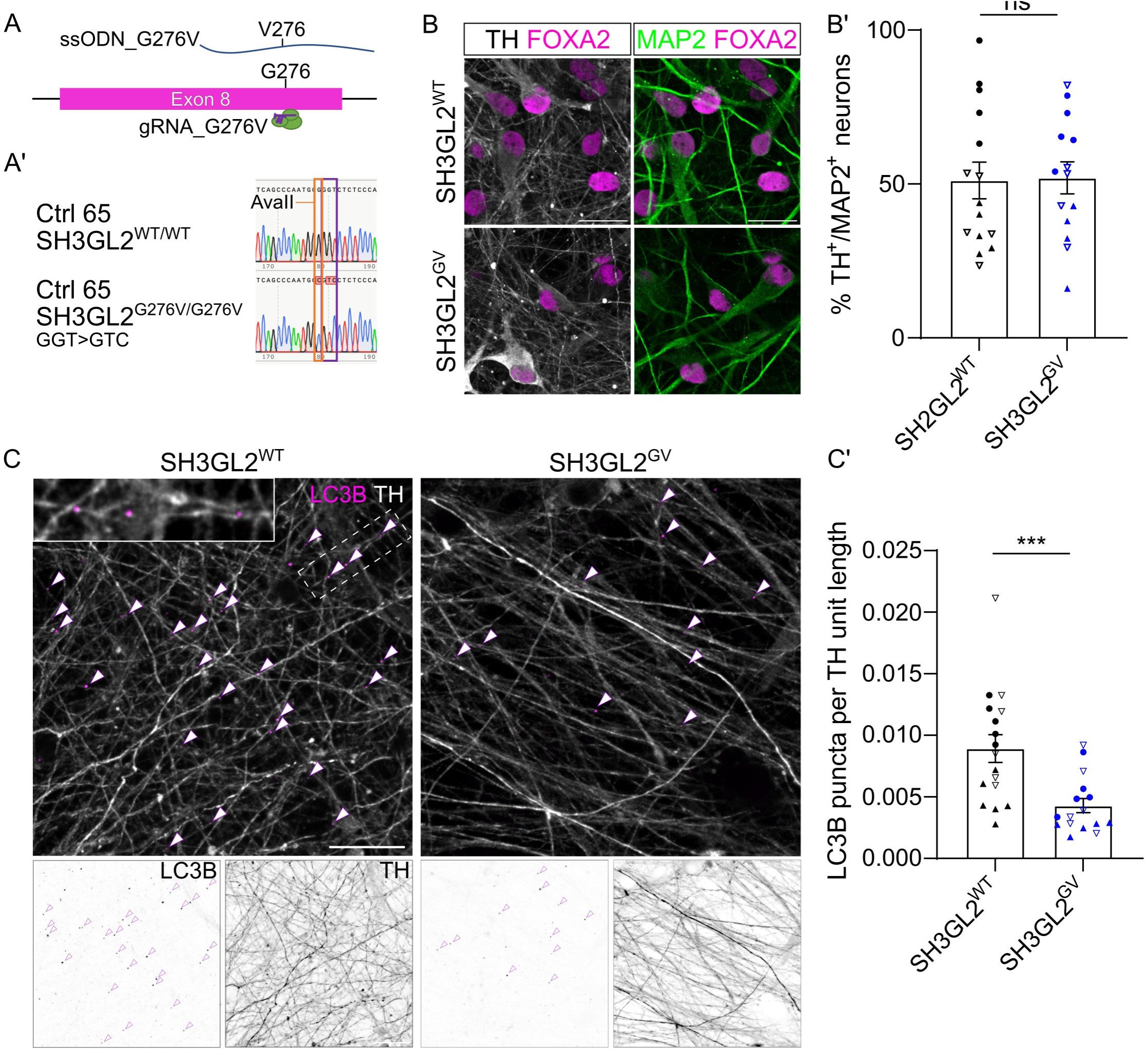
Expression of SH3GL2^G276V^ in differentiated dopaminergic neurons recapitulates findings from *Drosophila* synapses. (A) Schematic representation of gene editing strategy to knock-in the G276V mutation. (A’) Sanger sequencing of a single gene edited clone showing successful homozygouse editing of the indicated nucleotide. (B) Representative maximum projection confocal images of terminally differentiated (55-60 days) SH3GL2^WT^ vmDAn stained with the ventral midbrain marker FOXA2, dopaminergic marker TH and neuronal marker MAP2. (B’) Quantification of the amount of dopaminergic neurons over the total number of neurons in the field of view expressed as percentage of TH^+^/MAP2^+^ neurons. Scale bar: 100 µm. Error bars represent mean ± SEM. Statistical significance calculated with an ordinary one-way ANOVA with Tukey’s multiple comparison test: ns, not significant. Single data points representing single confocal images. Data from three independent vmDAn differentiations (circles and empty/filled triangles represent independent differentiations). (C) Representative maximum projection confocal images of terminally differentiated (55-60 days) SH3GL2^WT^ vmDAn stained with anti-TH and anti-LC3B antibodies. Arrowheads indicate LC3B^+^ autophagosomes within TH^+^ neurites. Scale bar: 20 µm. Insert box shows zoom of the region indicated within the dotted rectangle where multiple LC3B^+^ dots are visible. (C’) Quantification of the number of LC3B^+^ autophagosomes within TH^+^ neurites normalized on the total TH^+^ neurites length in the field of view. Error bars represent mean ± SEM. Statistical significance calculated with a student t-test. *\*\**\* *P* < 0.001. Single data points represent single confocal images. Data from three independent vmDAn differentiations (circles and empty/filled triangles represent independent differentiations).

## Discussion

This work reveals that EndoA links neuronal activity-dependent Ca^2+^ influx to synaptic autophagy and that a human mutation conferring risk to PD disrupts this process.

Synapses require regulated turnover systems to maintain protein homeostasis and disrupting this process causes synaptic and -ultimately- neuronal degeneration (Soukup et al., 2016; Vanhauwaert et al., 2017; Vijayan and Verstreken, 2017). There are several reasons why synapses are vulnerable, including that they need to cope with metabolically demanding processes, such as the synaptic vesicle cycle that is fueled by mitochondrial energy generation, thus causing the production of damaging ROS (Verstreken et al., 2005). Synapses are also often located at relatively long distances from neuronal cell bodies, imposing logistics problems that include the need to transport degradative organelles along long axons (Wang et al., 2020). Finally, individual synaptic boutons are fragile and uncontrolled autophagy could cause the (unwanted) engulfment of large parts of the cytoplasm, disrupting homeostasis. We propose that by coupling the activation of synaptic autophagy to neuronal activity, protein and organelle turnover are activated to maintain a healthy and functional synapse.

In this paper, we find that Ca^2+^ influx at stimulated synapses induces autophagosome formation. Work in AIY neurons in *C. elegans* also show a correlation between an increase in autophagosome formation and Ca^2+^ waves (Hill et al., 2019), suggesting this is a universal mechanism. The Ca^2+^ that induces synaptic autophagy upon stimulation appears to be largely derived from extracellular sources, as the L/N-type Ca^2+^ channel agonist Nefiracetam and the availability of extracellular Ca^2+^ promote the process (Nishizaki et al., 1998; Yoshii and Watabe, 1994; Yoshii et al., 2000). However, contributions from intracellular sources, such as ER-Ca^2+^, are not excluded or could further tune the process (Kuijpers et al., 2021) .

Mechanistically, we provide evidence that EndoA is critically required to relay this Ca^2+^ signal. *In vitro* work found a negatively charged residue at position 264 in the flexible region of rat ENDOA2 that is involved in regulating the Ca^2+^-dependent association of EndoA with binding partners at the plasma membrane, including Dynamin and voltage-gated Ca^2+^ channels (Chen et al., 2003; Zhang et al., 2012a; Kroll et al., 2019). We confirm this is the case for fly EndoA, but based on several biophysical methods, we do exclude large-scale Ca^2+^-induced conformational changes in EndoA. This is in agreement with recent data that also exclude direct binding of Ca^2+^ to EndoA (Yang et al., 2021) and leaves open the possibility that other, as of yet unidentified, intermediary partners could be needed. Nonetheless, our data clearly show that mutating the negatively charged residue of EndoA (D265 in flies) eliminates the protein’s proper response to Ca^2+^. Our data also indicate that Ca^2+^ regulates the mobility of EndoA in plasma-membrane nanodomains (this could be through binding the endocytic factor Dynamin) and its localization with nascent autophagosomes in the synapse lumen. Based on our results, we propose a model where under normal conditions, EndoA supports endocytic processes at the plasma membrane (Milosevic et al., 2011; Ringstad et al., 1999; Verstreken et al., 2002; Watanabe et al., 2018), and that Ca^2+^ influx at synapses liberates a (limited) pool of EndoA from nanodomains to drive synaptic autophagy. As only a fraction of EndoA re-localizes to the synapse lumen, leaving a large enough amount of protein at the membrane, both EndoA^D265R^ and EndoA^D265A^ can support endocytosis. The liberation of the EndoA-pool for autophagy appears a neuron- and synapse-specific process because KCl stimulation of chromaffin cells causes redistribution of EndoA towards the plasma membrane, suggesting a different mechanism is at play in these cells (Gowrisankaran et al., 2020).

The early stages of neurodegenerative diseases are associated with synaptic failure (Burke and O’Malley, 2013; Caminiti et al., 2017; Delva et al., 2020; Soukup et al., 2018). In PD, there is strong genetic evidence pointing to synaptic dysfunction and several proteins mutated in PD are enriched at synapses (eg. alpha-synuclein) and/or play essential roles in synaptic function (eg. DNAJC6 and Synj1) (Krebs et al., 2013; Krüger et al., 1998; Olgiati et al., 2016; Polymeropoulos et al., 1997; Proukakis et al., 2013; Quadri et al., 2013; Zarranz et al., 2004). Similarly, ENDOA1 is emerging as a key protein in PD and this is supported by human genetics: there are GWAS signals in the vicinity of the *SH3GL2* locus (Nalls et al., 2019) and there is the mutation associated with increased risk to PD in the open reading frame that we modelled here (Chang et al., 2017; Germer et al., 2019). We report that this PD risk mutation in ENDOA1 does not strongly affect synaptic vesicle endocytosis, but it does block synaptic autophagy. Excitingly, this role of EndoA in synaptic autophagy may be a central node in PD (Soukup et al., 2018) that includes Synj1 (that binds EndoA), LRRK2 (that phosphorylates EndoA) and possibly Parkin (that ubiquitinates EndoA) (Kitada et al., 1998; Krebs et al., 2013; Matta et al., 2012; Olgiati et al., 2014; Paisán-Ruiz, 2009). Mutations in each of these proteins causes alterations in autophagic function (Oliveira et al., 2015; Schöndorf et al., 2014; Soukup et al., 2016; Vanhauwaert et al., 2017; Yue et al., 2015) and also analyses of post-mortem tissue of PD patients found autophagy defects (Alvarez-Erviti et al., 2010; Murphy et al., 2015; Tanji et al., 2011). The exact function of the interactions between these PD-related proteins in pathology has not been elucidated. However, our finding that the PD risk mutant ENDOA1^G276V^ blocks stimulation-induced synaptic autophagy is an important piece in this puzzle and indicates the important role of synaptic autophagy in PD pathogenicity.

## STAR Methods

### Contact for Reagent and Resource Sharing

Further information and requests for resources and reagents should be directed to the Lead Contact Patrik Verstreken. Human control iPSC line (SFC065) was used in accordance with an MTA with the University of Lübeck (Germany).

### Experimental Model and Subject Details Fly stocks and maintenance

Flies were grown on standard cornmeal and sugar beet syrup medium at 25 °C. The *Drosophila* wild type and mutant (D265A, D265R) cDNA of EndophilinA, as well as human wild-type and mutant (G276V) SH3GL2 were cloned into the pUAST-attB vector and flies generated by in-house embryo injection or at Bestgene Inc into the VK37 locus to establish *UAS-EndoA^WT^, UAS-EndoA^D264A^, UAS-EndoA^D264R^, UAS-SH3GL2^WT^, UAS-SH3GL2^G276V^* lines and the mEOS3.1 tagged lines. Genomic *w^+^;;Atg8^3×mCherry^* (hereafter Atg8^mCherry^) flies were a gift from Gabor Juhasz (Hegedus et al., 2016).

### iPSC maintenance and differentiation

iPSC from a control line (SFC065) and *SH3GL2* p.G276V knock-in line were differentiated into ventral midbrain dopaminergic neurons as described in the methods section. Briefly, hiPSC were maintained on matrigel-coated plates with mTeSR-Plus medium (Stem Cell Technologies) and medium changes were performed every two days. At day 18 of differentiation, neural progenitors were plated on coverslips coated with poly-D-lysine and mouse laminin in terminal differentiation medium.

### Method details Genotypes

The following genotypes were used:

*w*;UAS-endoA^WT^/+;endoA^26^/endoA^Δ4^,GMR57C10-GAL4* (where *GMR57C10-Gal4* is referred to as *nSyb-Gal4*)

w*;UAS-endoA^D265A^/+;endoA^26^/endoA^Δ4^,GMR57C10-GAL4

w*;UAS-endoA^D265R^/+;endoA^26^/endoA^Δ4^,GMR57C10-GAL4

*w*;GMR57C10-GAL4/+;*

w*;; endoA^26^/endoA^Δ4^

_w_1118_CS_

*w^1118^;; genomic Atg8^3×mCherry^/+*

w*;genomic endoA^S75A^/+; endoA^26^/EndoA^Δ4^, genomic Atg8^3×mCherry^

*w*; UAS-RNAi-Atg3/GMR57C10-GAL4;genomic Atg8^3×mCherry^/+*

w*;UAS-endoA^WT^/GMR57C10-GAL4;endoA^26^/EndoA^Δ4^, genomic Atg8^3×mCherry^

w*;UAS-endoA^D265A^/GMR57C10-GAL4;endoA^26^/endoA^Δ4^, genomic Atg8^3×mCherry^

w*;UAS-endoA^D265R^/GMR57C10-GAL4;endoA^26^/endoA^Δ4^, genomic Atg8^3×mCherry^

w*;UAS-SH3GL2^WT^/GMR57C10-GAL4;endoA^26^/endoA^Δ4^, genomic Atg8^3×mCherry^

w*;UAS-SH3GL2^G276V^/GMR57C10-GAL4;endoA^26^/endoA^Δ4^, genomic Atg8^3×mCherry^

w*;UAS-SH3GL2^WT^/+;endoA^26^/endoA^Δ4^, GMR57C10-GAL4

*w*;UAS-SH3GL2^G276V^/+;endoA^26^/endoA^Δ4^, GMR57C10-GAL4w;GMR57C10-Gal4/+;*

*genomic Atg8^3×mCherry^/+*

*w*;UAS-GCaMP6m/+;GMR57C10-GAL4/+*

w*;UAS-endoA^WT^,cn bw/cn bw;endoA^26^/longGMR-GAL4

w*;UAS-endoA^D254A^,cn bw/cn bw;endoA^26^/longGMR-GAL4

w*;UAS-endoA^D254R^,cn bw/cn bw;endoA^26^/longGMR-GAL4

*w*;cn bw/cn bw;longGMR-GAL4/+*

w*;UAS-mEOS3.1::endoA^WT^/+;endoA^26^/endoA^Δ4^,GMR57C10-GAL4

w*;UAS-mEOS3.1::endoA^D265A^/+;endoA^26^/endoA^Δ4^,GMR57C10-GAL4

w*;UAS-mEOS3.1::endoA^D265R^/+;endoA^26^/endoA^Δ4^,GMR57C10-GAL4

### Primers, gBlocks and Oligos

- pUAST_EndoA-WT

FW EndoA WT for UAS: ACTCTGAATAGGGAATTGGGATGGCTTTCGCCGGACTCAA AAAGC

Rc EndoA WT for UAS: AAAGATCCTCTAGAGGTACCCCTAGTTGCCATTGGGCAGG Gibson Assembly with EndoA PCR fragment and pUAST_attB linearized with EcoRI and XhoI

- pUAST_EndoA-D265R

EndoA D265R gBlock

ACTCTGAATAGGGAATTGGGCAAAATGGCTTTCGCCGGACTCAAAAAGCAGATCAACAAGGCCAACCAGTATATGACGGAGAAGATGGGCGGTGCCGAGGGCACCAAACTGGACATGGACTTCATGGAGATGGAACGCAAGACGGACGTCACCGTGGAGCTAGTGGAGGAGCTGCAGCTAAAGACGAAGGAGTTCCTGCAGCCGAATCCAACGGCACGGGCCAAAATGGCAGCGGTCAAGGGCATCTCGAAGCTGTCCGGACAGGCCAAGTCCAATACGTATCCGCAACCGGAGGGCCTGCTCGCGGAATGCATGCTGACTTATGGGAAGAAGCTCGGCGAGGACAACAGCGTGTTCGCGCAGGCGCTCGTCGAATTCGGCGAAGCGCTGAAACAGATGGCCGACGTCAAGTATTCGCTGGACGACAACATCAAGCAGAACTTTTTGGAGCCACTGCATCATATGCAGACCAAAGACCTCAAGGAGGTAATGCATCATCGCAAGAAGCTGCAGGGCCGGCGGCTAGACTTTGACTGCAAGCGTCGCCGACAGGCCAAGGACGATGAGATTCGTGGTGCCGAGGACAAGTTCGGTGAATCGCTCCAGCTGGCCCAGGTGGGCATGTTCAATTTGCTCGAGAACGATACGGAGCATGTCTCCCAGCTGGTCACCTTTGCCGAGGCACTATACGATTTTCATTCGCAATGCGCGGATGTCCTTCGAGGCCTGCAGGAGACACTGCAGGAGAAGCGCTCCGAGGCGGAGAGCCGGCCACGCAACGAGTTCGTGCCCAAGACGCTGCTCGATCTGAACTTGcgCGGCGGTGGCGGCGGCCTCAACGAAGATGGCACGCCGTCTCACATTAGTTCGAGCGCCTCGCCGTTGCCCTCGCCGATGCGTTCGCCCGCCAAGTCGATGGCCGTAACGCCGCAGCGCCAGCAGCAGCCCTGCTGCCAGGCCCTCTACGACTTCGAGCCGGAGAATCCCGGCGAACTGGCCTTCAAGGAGAACGACATTATCACCCTGTTGAATCGCGTCGACGACAATTGGTTCGAGGGCGCGGTGAATGGCCGCACCGGTTACTTCCCGCAGTCGTATGTTCAGGTGCAGGTGCCCCTGCCCAATGGCAACTAGGGGTACCTCTAGAGGATCTTT

Gibson Assembly with EndoA D265R gBlock and pUAST_attB linearized with EcoRI and XhoI

- pUAST_EndoA-D265A

EndoA D264A gBlock

ACTCTGAATAGGGAATTGGGATGGCTTTCGCCGGACTCAAAAAGCAGATCAACAAGGCCAACCAGTATATGACGGAGAAGATGGGCGGTGCCGAGGGCACCAAACTGGACATGGACTTCATGGAGATGGAACGCAAGACGGACGTCACCGTGGAGCTAGTGGAGGAGCTGCAGCTAAAGACGAAGGAGTTCCTGCAGCCGAATCCAACGGCACGGGCCAAAATGGCAGCGGTCAAGGGCATCTCGAAGCTGTCCGGACAGGCCAAGTCCAATACGTATCCGCAACCGGAGGGCCTGCTCGCGGAATGCATGCTGACTTATGGGAAGAAGCTCGGCGAGGACAACAGCGTGTTCGCGCAGGCGCTCGTCGAATTCGGCGAAGCGCTGAAACAGATGGCCGACGTCAAGTATTCGCTGGACGACAACATCAAGCAGAACTTTTTGGAGCCACTGCATCATATGCAGACCAAAGACCTCAAGGAGGTAATGCATCATCGCAAGAAGCTGCAGGGCCGGCGGCTAGACTTTGACTGCAAGCGTCGCCGACAGGCCAAGGACGATGAGATTCGTGGTGCCGAGGACAAGTTCGGTGAATCGCTCCAGCTGGCCCAGGTGGGCATGTTCAATTTGCTCGAGAACGATACGGAGCATGTCTCCCAGCTGGTCACCTTTGCCGAGGCACTATACGATTTTCATTCGCAATGCGCGGATGTCCTTCGAGGCCTGCAGGAGACACTGCAGGAGAAGCGCTCCGAGGCGGAGAGCCGGCCACGCAACGAGTTCGTGCCCAAGACGCTGCTCGATCTGAACTTGGcCGGCGGTGGCGGCGGCCTCAACGAAGATGGCACGCCGTCTCACATTAGTTCGAGCGCCTCGCCGTTGCCCTCGCCGATGCGTTCGCCCGCCAAGTCGATGGCCGTAACGCCGCAGCGCCAGCAGCAGCCCTGCTGCCAGGCCCTCTACGACTTCGAGCCGGAGAATCCCGGCGAACTGGCCTTCAAGGAGAACGACATTATCACCCTGTTGAATCGCGTCGACGACAATTGGTTCGAGGGCGCGGTGAATGGCCGCACCGGTTACTTCCCGCAGTCGTATGTTCAGGTGCAGGTGCCCCTGCCCAATGGCAACTAGGGGTACCTCTAGAGGATCTTT

Gibson Assembly with EndoA D265A gBlock and pUAST_attB linearized with EcoRI and XhoI pUAST_EndoA-WT-mEOS3.1-v5, pUAST_EndoA-D265R-mEOS3.1-v5, pUAST_EndoA-D265A-mEOS3.1-v5 were made by respectively performing a restriction digest on pUAST_EndoA-WT, pUAST_EndoA-D264R, pUAST_EndoA-D265A with AgeI and XbaI and doing a Gibson assembly with:

gBlock mEOS3.1

GAGGGCGCGGTGAATGGCCGCACCGGTTACTTCCCGCAaTCGTATGTTCAGGTGCAGGTtCCCCTtCCCAATGGCAACGGTGGGGGTACgGGAGGCGGATCCATGAGTGCGATTAAGCCAGACATGAAGATCAAACTCCGTATGGAAGGCAACGTAAACGGGCACCACTTTGTGATCGACGGAGATGGTACAGGCAAGCCTTTTGAGGGAAAACAGAGTATGGATCTTGAAGTCAAAGAGGGCGGACCTCTGCCTTTTGCCTTTGATATCCTGACCACcGCATTCCATTACGGCAACAGGGTATTCGCCAAATATCCAGACAACATACAAGACTATTTTAAGCAGTCGTTTCCTAAGGGGTATTCGTGGGAACGAAGCTTGACTTTCGAAGACGGGGGCATTTGCAACGCCAGAAACGACATAACAATGGAAGGGGACACTTTCTATAATAAAGTTCGATTTTATGGTACCAACTTTCCCGCCAATGGTCCAGTTATGCAGAAGAAGACGCTGAAATGGGAGCCCTCCACTGAGAAAATGTATGTGCGTGATGGAGTGCTGACGGGTGATATTGAGATGGCTTTGTTGCTTGAAGGAAATGCCCATTACCGATGTGACTTCAGAACTACTTACAAAGCTAAGGAGAAGGGTGTCAAGTTACCAGGCGCCCACTTTGTGGACCACTGCATTGAGATTTTAAGCCATGACAAAGATTACAACAAGGTTAAGCTGTATGAGCATGCTGTTGCTCATTCTGGATTGCCTGACAATGCCAGACGAGGaGGaGGTACCGGAGGtGGcTCCGGCAAGCCCATCCCCAACCCCCTGCTGGGCCTGGATAGCACCTAGAGGATCTTTGTGAAGGAACCTTAC

All plasmids contain the EndoA CDS (WT or mutation), a short flexible linker (GGTGGS), mEOS3.1, again a short flexible linker (GGTGGS) and a V5 epitope tag.

- pUAST_SH3GL2-WT

SH3GL2 WT gBlock

AACTCTGAATAGGGAATTGGGCAAAAATGTCGGTGGCCGGCCTCAAGAAGCAGTTCCATAAAGCCACTCAGAAAGTGAGTGAGAAGGTTGGAGGAGCTGAAGGAACCAAGCTAGATGATGACTTCAAAGAGATGGAAAGGAAAGTGGATGTCACCAGCAGGGCTGTGATGGAAATAATGACTAAAACAATTGAATACCTTCAACCCAATCCAGCTTCCAGAGCTAAGCTCAGCATGATCAACACCATGTCAAAAATCCGTGGCCAGGAGAAGGGGCCAGGCTATCCTCAGGCAGAGGCGCTGCTGGCAGAGGCCATGCTCAAATTTGGAAGAGAGCTTGGAGATGATTGCAACTTTGGCCCAGCACTTGGTGAGGTCGGGGAGGCCATGCGGGAACTGTCGGAGGTCAAAGACTCTTTGGACATAGAAGTGAAGCAGAACTTCATTGACCCTCTTCAGAATCTTCATGACAAAGATCTTAGGGAAATTCAACATCATCTAAAGAAGTTGGAGGGTCGACGCCTGGATTTTGATTATAAGAAGAAACGACAAGGCAAGATTCCGGATGAAGAGCTTCGTCAAGCTCTAGAGAAATTTGATGAGTCTAAGGAAATTGCTGAGTCAAGCATGTTCAATCTCTTGGAGATGGATATTGAACAAGTGAGCCAGCTCTCTGCACTTGTGCAAGCTCAGCTGGAGTACCACAAGCAGGCAGTCCAGATCCTGCAGCAAGTCACGGTCAGACTGGAAGAAAGAATAAGACAGGCTTCATCTCAGCCTAGAAGGGAATATCAACCTAAACCACGAATGAGCCTGGAGTTTCCAACTGGAGACAGTACTCAGCCCAATGGGGGTCTCTCCCACACAGGCACTCCCAAACCTTCAGGTGTCCAAATGGATCAGCCgTGCTGCCGAGCTCTGTACGACTTTGAACCTGAAAATGAAGGGGAGTTGGGATTTAAAGAGGGCGATATCATCACACTCACTAACCAAATTGATGAGAACTGGTATGAGGGGATGCTGCATGGCCATTCAGGCTTCTTCCCCATCAATTATGTGGAAATTCTGGTTGCCCTGCCCCATTAGGGGTACCTCTAGAGGATCTTTG

Gibson Assembly with SH3GL2 WT gBlock and pUAST_attB linearized with EcoRI and XhoI

- pUAST_SH3GL2-G276V SH3GL2 G276V gBlock

AACTCTGAATAGGGAATTGGGCAAAAATGTCGGTGGCCGGCCTCAAGAAGCAGTTCCATAAAGCCACTCAGAAAGTGAGTGAGAAGGTTGGAGGAGCTGAAGGAACCAAGCTAGATGATGACTTCAAAGAGATGGAAAGGAAAGTGGATGTCACCAGCAGGGCTGTGATGGAAATAATGACTAAAACAATTGAATACCTTCAACCCAATCCAGCTTCCAGAGCTAAGCTCAGCATGATCAACACCATGTCAAAAATCCGTGGCCAGGAGAAGGGGCCAGGCTATCCTCAGGCAGAGGCGCTGCTGGCAGAGGCCATGCTCAAATTTGGAAGAGAGCTTGGAGATGATTGCAACTTTGGCCCAGCACTTGGTGAGGTCGGGGAGGCCATGCGGGAACTGTCGGAGGTCAAAGACTCTTTGGACATAGAAGTGAAGCAGAACTTCATTGACCCTCTTCAGAATCTTCATGACAAAGATCTTAGGGAAATTCAACATCATCTAAAGAAGTTGGAGGGTCGACGCCTGGATTTTGATTATAAGAAGAAACGACAAGGCAAGATTCCGGATGAAGAGCTTCGTCAAGCTCTAGAGAAATTTGATGAGTCTAAGGAAATTGCTGAGTCAAGCATGTTCAATCTCTTGGAGATGGATATTGAACAAGTGAGCCAGCTCTCTGCACTTGTGCAAGCTCAGCTGGAGTACCACAAGCAGGCAGTCCAGATCCTGCAGCAAGTCACGGTCAGACTGGAAGAAAGAATAAGACAGGCTTCATCTCAGCCTAGAAGGGAATATCAACCTAAACCACGAATGAGCCTGGAGTTTCCAACTGGAGACAGTACTCAGCCCAATGGGGtTCTCTCCCACACAGGCACTCCCAAACCTTCAGGTGTCCAAATGGATCAGCCgTGCTGCCGAGCTCTGTACGACTTTGAACCTGAAAATGAAGGGGAGTTGGGATTTAAAGAGGGCGATATCATCACACTCACTAACCAAATTGATGAGAACTGGTATGAGGGGATGCTGCATGGCCATTCAGGCTTCTTCCCCATCAATTATGTGGAAATTCTGGTTGCCCTGCCCCATTAGGGGTACCTCTAGAGGATCTTTG

Gibson Assembly with SH3GL2 G276V gBlock and pUAST_attB linearized with EcoRI and XhoI

pGEX-6P-1_EndoA-WT, pGEX-6P-1_EndoA-D265A and pGEX-6P-1_EndoA-D265R were generated by performing a PCR respectively from pUAST_EndoA-WT, pUAST_EndoA-D265A and pUAST_EndoA-D265R with the following primers:

FW: TTCTGTTCCAGGGGCCCCTGGGATCCATGGCTTTCGCCGGACTC Rc: GCGGCCGCTCGAGTCGACCCGGGCTAGTTGCCATTGGGCAG

Gibson assembly was performed with the PCR fragments and pGEX-6P-1 linearized with BamHI and EcoRI

- pGEX-6P-1_SH3GL2-WT

SH3GL2 WT gBlock: the sequence was codon optimized for expression in E.coli

TTCTGTTCCAGGGGCCCCTGGGATCCATGAGCGTCGCAGGCCTGAAGAAGCAGTTCCATAAGGCTACTCAGAAGGTTTCGGAAAAAGTAGGTGGGGCTGAGGGCACCAAGTTAGACGACGACTTTAAGGAAATGGAAAGAAAAGTCGATGTCACGAGTAGAGCGGTTATGGAAATTATGACGAAGACCATAGAGTATTTGCAGCCGAATCCCGCCAGTCGTGCCAAGTTGAGCATGATCAATACGATGTCGAAAATTCGCGGGCAGGAAAAAGGACCGGGATACCCTCAAGCAGAGGCGCTTCTTGCAGAGGCCATGTTAAAATTTGGGCGCGAGCTTGGAGATGACTGCAATTTTGGCCCAGCTTTAGGGGAGGTTGGTGAGGCAATGAGAGAGTTATCCGAAGTCAAGGATTCCCTGGACATTGAGGTTAAGCAGAACTTTATAGATCCACTTCAAAATTTGCACGATAAAGACCTGCGTGAGATTCAACATCACTTAAAGAAGTTAGAAGGACGCCGCCTTGACTTTGATTATAAGAAAAAGCGTCAGGGCAAAATACCCGACGAAGAACTTCGCCAAGCTCTGGAGAAGTTTGATGAGAGCAAGGAAATAGCTGAAAGTTCGATGTTTAATTTGCTGGAGATGGATATTGAACAAGTAAGTCAGTTATCTGCGTTAGTACAGGCCCAATTAGAATATCACAAACAGGCTGTGCAAATATTACAACAGGTAACCGTACGTTTGGAGGAGAGAATACGTCAGGCATCGTCTCAGCCTCGCCGCGAATACCAACCGAAACCACGCATGTCGCTTGAGTTCCCCACAGGAGACTCAACCCAACCTAACGGAGGCTTGTCACATACGGGCACACCCAAGCCGAGCGGAGTTCAGATGGACCAGCCGTGCTGTAGAGCACTGTATGATTTCGAGCCGGAGAACGAGGGTGAGCTTGGGTTCAAGGAGGGGGATATTATTACTCTTACTAATCAGATTGATGAGAATTGGTACGAGGGGATGCTTCATGGTCATTCGGGCTTTTTCCCTATAAATTACGTCGAGATTCTGGTGGCGCTGCCACACTAGCCCGGGTCGACTCGAGCGGCCGCATCGTGACTGACTGAC

Gibson assembly performed with codon optimized SH3GL2 WT gBlock and pGEX-6P-1 linearized with BamHI and EcoRI

- pGEX-6P-1_SH3GL2-G267V

SH3GL2 G276V gBlock: the sequence was codon optimized for expression in E.coli

TTCTGTTCCAGGGGCCCCTGGGATCCATGAGCGTCGCAGGCCTGAAGAAGCAGTTCCATAAGGCTACTCAGAAGGTTTCGGAAAAAGTAGGTGGGGCTGAGGGCACCAAGTTAGACGACGACTTTAAGGAAATGGAAAGAAAAGTCGATGTCACGAGTAGAGCGGTTATGGAAATTATGACGAAGACCATAGAGTATTTGCAGCCGAATCCCGCCAGTCGTGCCAAGTTGAGCATGATCAATACGATGTCGAAAATTCGCGGGCAGGAAAAAGGACCGGGATACCCTCAAGCAGAGGCGCTTCTTGCAGAGGCCATGTTAAAATTTGGGCGCGAGCTTGGAGATGACTGCAATTTTGGCCCAGCTTTAGGGGAGGTTGGTGAGGCAATGAGAGAGTTATCCGAAGTCAAGGATTCCCTGGACATTGAGGTTAAGCA GAACTTTATAGATCCACTTCAAAATTTGCACGATAAAGACCTGCGTGAGATTCAACATCACTTAAAGAAGTTAGAAGGACGCCGCCTTGACTTTGATTATAAGAAAAAGCGTCAGGGCAAAATACCCGACGAAGAACTTCGCCAAGCTCTGGAGAAGTTTGATGAGAGCAAGGAAATAGCTGAAAGTTCGATGTTTAATTTGCTGGAGATGGATATTGAACAAGTAAGTCAGTTATCTGCGTTAGTACAGGCCCAATTAGAATATCACAAACAGGCTGTGCAAATATTACAACAGGTAACCGTACGTTTGGAGGAGAGAATACGTCAGGCATCGTCTCAGCCTCGCCGCGAATACCAACCGAAACCACGCATGTCGCTTGAGTTCCCCACAGGAGACTCAACCCAACCTAACGGAGTCTTGTCACATACGGGCACACCCAAGCCGAGCGGAGTTCAGATGGACCAGCCGTGCTGTAGAGCACTGTATGATTTCGAGCCGGAGAACGAGGGTGAGCTTGGGTTCAAGGAGGGGGATATTATTACTCTTACTAATCAGATTGATGAGAATTGGTACGAGGGGATGCTTCATGGTCATTCGGGCTTTTTCCCTATAAATTACGTCGAGATTCTGGTGGCGCTGCCACACTAGCCCGGGTCGACTCGAGCGGCCGCATCGTGACTGACTGAC

Gibson assembly performed with codon optimized SH3GL2 G276V gBlock and pGEX-6P-1 linearized with BamHI and EcoRI

To generate G276V knock-in iPSCs the following Oligos and gRNAs were used:

ssODN_G276V_AvaII: cgaatgagcctgGAGtttccaactggagacagtactcagcccaatggCgtCctctcccacacaggcactcccaaaccttcaggtaagagctgaaactgca

px_SH3GL2_G276V_gRNA1_Fwd: CACCGTGTGTGGGAGAGACCCCCAT

px_SH3GL2_G276V_gRNA1_Rv: aaacATGGGGGTCTCTCCCACACAc

SH3GL2(G276V)_pRR_Fwd: tatcaacctaaaccacgaatgagcctggagtttccaactggagACGT

SH3GL2(G276V)_pRR_Rv: TCAGCCCAATGGGGGTCTCTCCCACACAGGCACTAGCT

PCR validation of the edited clones were carried out with the following primers:

SH3GL2_V-sp_Rv: gcctgtgtgggagagGacg

SH3GL2_Exon8_Rv: agtttctacctgacaactgactcc

SH3GL2_T7_Fwd: catggtagcatggtgggtgac

### Autophagy induction assays

Third instar larvae (still crawling in the food) expressing Atg8^mCherry^ were dissected in fresh Ca^2+^-free HL3 (100 mM NaCl, 5 mM KCl, 10 mM NaHCO_3_, 5 mM Hepes, 30 mM sucrose, 5 mM trehalose and 10 mM MgCl_2_, pH 7.2).

For electrical stimulation the nerves of dissected larvae were cut just below the ventral nerve cord, and axons innervating segments A3 and A4 were stimulated at 20 Hz for 30 min at 50 % above the threshold using a suction electrode (Soukup et al., 2016). During electrical stimulation, larvae were incubated in HL3 supplemented with or without 1 mM CaCl_2_, as indicated. EGTA-AM (Thermo Fisher) was used at 25 µM (in anhydrous 0.1 % DMSO).

For calcium channel agonist treatment larvae were prepared as for electrical stimulation and then incubated for 30 min in HL3 with 10 µM of Nefiracetam (Tocris), dissolved in 0,0025 % anhydrous DMSO (Yoshii and Watabe, 1994; Yoshii et al., 2000), and with or without 1 mM CaCl_2_. 100 µM of NAS was similarly used to prevent muscle contractions.

In all experiments 100 µM 1-Naphthylacetyl spermine trihydrochloride – NAS (Sigma) was added (Levitan et al., 2007; Soukup et al., 2016) to prevent muscle contractions.

### Confocal Live Imaging and quantification

Live imaging of dissected larvae was carried out on a Nikon A1R confocal microscope with a 60X (NA 1.0) water-dipping lens. Dissected larvae were washed multiple times in HL3 prior to imaging. NIS Elements (Nikon) was used for data acquisition using the Resonant Scanning option, with a zoom factor of 3 and line averaging of 16. All images were acquired with a pinhole of one Airy unit and a resolution of 1024 × 1024. Z-stacks were used in data acquisition to capture fluorescence (puncta presence) across the whole NMJ. Low basal autophagy levels were always confirmed in control experiments.

Quantification of the number of puncta within synaptic boutons was performed in ImageJ (Soukup et al., 2016). Atg8^3×mCherry^ puncta were manually counted and this was combined with the application of a threshold mask as to determine the area of the bouton. Trafficking autophagosomes (punctae in motor neuron axons outside the NMJ) were excluded from data analysis. The conditions were blinded for quantification.

### Calcium imaging

Third instar larvae expressing GCaMP6m in neurons (GMR57C10-GAL4) were dissected in fresh HL3 and nerves cut below the ventral nerve cord. After dissection, HL3 was replaced with freshly prepared 100 µM Nas (Sigma) and 2 mM CaCl_2_ in HL3. Larval filets were imaged on an widefield Nikon Eclipse FN1 upright microscope equipped with a water dipping 20X lens (NA 0.95) and a GFP filter. Time-lapse images were acquired at a frequency of 5 Hz. Nefiracetam or DMSO (control) was added during imaging. Image analysis was performed using ImageJ. GCaMP6m intensity was plotted as ΔF/F_0_ and measured, after background subtraction, by normalizing the signal, within the synaptic boutons, in each frame to the averaged GCaMP6m intensity, for the same region of interest, of the first 10 frames.

### Immunohistochemistry and confocal imaging

Third instar larvae were dissected in cold Ca^2+^ free HL3 and fixed for 20 minutes at room temperature with 4 % para-formaldehyde. Fixed larvae were permeabilized with 0.4 % PBX (TritonX-100 in 1X PBS), blocked for 1 hour with 10 % normal goat serum in PBX and incubated overnight at 4°C with primary antibodies. After several washes, larval filets were incubated with secondary antibodies for 90 min at room temperature. Samples were mounted in Vectashield (Vector Laboratories).

The following antibodies were used: guinea pig anti-EndoA (GP69) [1:2000 (Verstreken et al., 2002)], rabbit anti-HRP [1:1000 (Jackson ImmunoReasearch)], anti-SH3GL2 [1:1000 (GeneTex)]. Alexa Fluor-488/Alexa Fluor-555 conjugated secondary antibodies [1:1000 (Invitrogen)].

Samples were imaged on a Nikon A1R confocal microscope with a 60X (NA 1.4) oil lens. Acquisition performed using a resonant scanner, a zoom factor of 3 and line averaging of 16. All images were acquired with a pinhole of 1 Airy unit and a resolution of 1024 × 1024. Z-stacks (step intervals of 0.45 µm) were used in data acquisition and the same image settings were maintained across the genotypes. Quantification of fluorescent intensity and NMJ area was performed with ImageJ.

For the characterization of iPSCs and midbrain floor plate progenitors, cells were fixed at day 16 of differentiation for 15 min in 4 % para-formaldehyde. Cells were blocked for 1 h at room temperature with 3 % normal goat serum + 0.3 % Triton X-100 (Sigma) in DPBS supplemented with Ca^2+^ and Mg^2+^ (Life Technologies). Primary antibodies were incubated overnight at 4°C in blocking solution. Secondary antibodies were incubated for 1 h at room temperature in blocking solution. Coverslips were mounted in Mowiol (Sigma) and imaged on an upright Nikon A1R confocal microscope equipped with a DIC N2 20X lens (NA 0.75). Z-stacks were acquired with pinhole of 1 Airy unit, a Galvano scanner with line averaging of 2, image size of 1024 x 1024 pixels and step intervals of 2 µm (for imaging of iPSCs) and 0.5 µm (for imaging of midbrain floor plate progenitors).

For the quantification of autophagy induction, terminally differentiated vmDAn (55-60 days) were fixed for 15 min in 4 % para-formaldehyde and blocked for 1 h at room temperature in 3 % normal goat serum + 0.01 % Saponin (Sigma) in DPBS supplemented with with Ca^2+^ and Mg^2+^ (Life Technologies). Primary antibodies were diluted in blocking solution and incubated overnight at 4°C, while secondary antibodies were incubated for 1 h at room temperature. Coverslips were mounted with Mowiol (Sigma) and imaged on an upright Nikon A1R confocal microscope equipped with an oil immersion Apo 60X lens with (NA 1.4). Z-stacks were acquired with a pinhole of 1 Airy unit, a Galvano scanner with line averaging of 2, image size of 1024 x 1024 pixels and step intervals of 0.3 µm.

The following antibodies were used: mouse IgG1 anti-SOX2 [1:200 (Santa Cruz)], rabbit anti-OCT4 [1:50 (Abcam)], mouse IgG1 anti-NANOG [1:50 (Santa Cruz)], rabbit anti-LMX1A/B [1:1000 (Millipore)], mouse IgG2a anti-FOXA2 [1:250 (Santa Cruz)], mouse IgG1 anti-Engrailed-1 [1:100 (DSHB, 4G11)], mouse IgG2a anti-OTX2 [1:100 (Santa Cruz)], Rabbit anti-LC3B [1:250 (Cell Signaling)], mouse IgG2a anti-TH [1:500 (Santa Cruz)], Chicken anti-MAP2 [1:500 (Abcam)], mouse IgG1 anti-alpha-Tubulin [1:500 (DSHB 12G10)], Alexa Fluor-488/ Alexa Fluor-555/ Alexa Fluor-647 conjugated secondary antibodies [1:500 (Invitrogen)].

### Zeiss Airyscan Confocal Imaging and quantification

A Zeiss LSM 880 (AiryScan detector enabled) with a 63X (NA 1.4) was used to image the distribution of EndoA (WT, D264A and D264R) across the NMJ. Zen Black software (2012, Carl Zeiss) was used for image acquisition and processing.

To quantify the distribution of EndoA within boutons, we selected boutons of similar size as previously described (Kasprowicz et al., 2014). These were then rescaled to a standard diameter and average the labelling intensities per position along the bouton diameter were defined. Boutons were resized to a width of 500 pixels, and integrated intensity was calculated for whole boutons fit into a 400 pixel diameter, while cytosol integrated intensity was calculated by 250 pixel diameter circle (in ImageJ). The average integrated intensity was calculated across all boutons of the same genotype.

### Single-particle tracking photoactivated localization microscopy (sptPALM)

SptPALM was carried out on transgenic EndophilinA-mEOS3.1 (wild-type, D264A and D264R) expressing *Drosophila* third-instar larvae which were dissected on a PDMS (Sylgard) base as previously described (Bademosi et al., 2017, 2018; Vanhauwaert et al., 2017). Briefly, EndoA^WT::mEos3.1^, EndoA^DA:mEos3.1^ and EndoA^DR:mEos3.1^ were imaged at the larvae synaptic boutons on muscle 13 of segments A3 and A4 using total internal reflection (TIRF) microscopy under slightly oblique illumination. The dissected larvae were inverted onto glass-bottomed imaging dishes (MatTek Corporation). The larvae were perfused with HL3, or a solution mixture of HL3, 10 μM Nefiracetam, 100 µM NAS and 1 mM CaCl_2_. Acquisition of single EndoA-mEOS3.1 molecules was carried out using a C-Apochromat 63X (NA 1.2) water objective on the ELYRA PS.1 microscope (Zeiss). Synaptic boutons were located using 488 nm laser illumination. A 405 nm laser was used for photoconversion, and the 561 nm laser was used during acquisition. To spatially visualize and temporally characterize individual photoconverted fluorophores, the 405 nm laser was used at 0.00003-0.03 % power, while the 561 nm laser was used at 20 % power. A sensitive electron-multiplying charge-coupled device (EMCCD) camera (Evolve, Photomertic) was used to collect single molecule fluorescence. Zen Black acquisition software (2012 version, Carl Zeiss) was utilized for movie acquisition. 15,000 frames images were captured per synaptic bouton at a frame capture rate of 20 Hz. Analysis of sptPALM movies is described in the Supplemental Experimental Procedure.

### sptPALM Analysis

NIH ImageJ was used to convert movies from Zen’s CZI format to the format recognizable for the image analysis software – Tiff format. PALM-Tracer, a customized-written software plugin in Metamorph (Molecular Devices) (Kechkar et al., 2013; Nair et al., 2013) was used to localize single molecules and quantify their mobility. Mobility data was plotted as mean square displacement (µm^2^) values (equation 1) as well as diffusion coefficient (µm^2^ s^-1^) values (equation 2). The parameters were set to isolate and recognize trajectories of molecules undergoing free cytosolic diffusion, associated with organelles or bound to presynaptic plasma membrane. Only fluorescent molecules with sufficient threshold and with consecutive appearance across eight movie frames were tracked and used for data analysis. The PALM-Tracer software generated trajectory maps, super-resolved average intensity as well as diffusion coefficient maps. The colour gradient in the trajectory maps indicate the time of detection of the tracks; the blue trajectories indicate molecules detected early during movie acquisition, while the white trajectories indicate later appearance and detection.

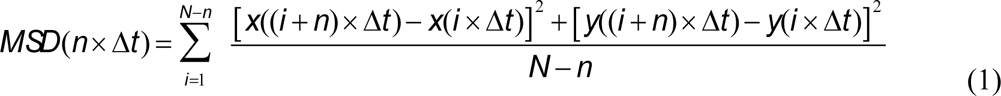

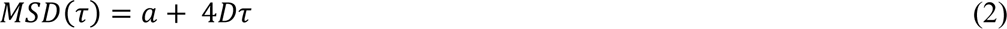

N is the number of data points in a trajectory, *Δt* is the time interval of each frame, *x* and *y* are the coordinates of a particle, a is the offset constant which incorporates the effects of localization error and finite camera exposure, and *D* is the diffusion coefficient. The MSD was calculated for the time interval *τ = nΔt* for the entire duration of each trajectory. The first eight points of the MSD were averaged over all trajectories and plotted against time.

The diffusion coefficient, *D*, (µm^2^/second) was calculated for each trajectory, from linear fits of the first four points of the MSD versus time function using equation. The diffusion coefficient (D) distribution was sorted into two populations, immobile and mobile. The immobile population of molecules explore an area inferior to that defined by the spatial resolution within one frame. The D_threshold_ = 0.0316 μm^2^/s was calculated as previously described (Constals et al., 2015). The immobile population was composed of molecules with a D value lower than 0.0316 μm^2^/s, while the mobile population was composed of molecules with D values above 0.0316 μm^2^/s.

The Robust regression and Outlier (ROUT) outlier test on Graph Pad Prism was used to identify and remove outliers. Relative frequency distribution graphs and average MSD curves were obtained using Graph Prism (version 6.0).

### Single-Molecule Localization Microscopy and Cluster Analysis

EndoA^::mEos3.1^ expressing larvae were dissected in either HL3 with Nefiracetam and NAS with or without 1 mM CaCl_2_ and then fixed with 4 % para-formaldehyde. Single-molecule localization was carried out on an ELYRA PS.1 microscope (Zeiss). Each dataset was acquired at a rate of 20 Hz for a duration of 20,000 frames by which point the mEOS3.1-tagged molecule were fully photoconverted. Coordinates of individual localizations were retrieved from each time-lapse video using Zen software (Zeiss). The datasets were corrected for *x,y* drift using Zen’s automated fiducial markers and affline transform algorithms. Localizations that appeared within 1 frame and 1 pixel of each other were consolidated to account for individual fluorophore blinking. Density-based spatial clustering of applications with noise (DBSCAN) analyses, a spatial clustering algorithm based on density, was used to quantify the clustering of proteins (Ester et al., 1996). DBSCAN identifies clusters in large datasets of localization coordinates by a continuous and propagative method that links components of a common cluster based on two parameters, r, and ε. Were r is the search radius and ε is the minimum number of neighbouring localizations within the search radius. Localizations outside the search radius and neighbouring points were classified as noise. The DBSCAN was implemented in Python.

### Transmission Electron Microscopy (TEM)

Third instar larvae were dissected in cold HL3 and processed for transmission electron microscopy as previously described (Lauwers et al., 2018). Briefly, larval fillets were fixed in fresh 4 % para-formaldehyde (Electron Microscopy Sciences) and 1 % glutaraldehyde (Sigma) in 1 mM MgCl_2_ (Sigma) and 0.1 M Na-cacodylate (Sigma) buffer, pH 7.2, overnight at 4°C. Samples were washed with 0.1 M Na-cacodylate, pH 7.4 and osmicated with 2 % osmium (OsO_4_/Na-Cacodylate buffer). Next, tissue was stained with 2 % uranyl acetate (Electron Microscopy Sciences) for 1.5 h and embedded in Agar 100 resin (Agar Scientific). Horizontal ultrathin sections (70 nm) were collected. Synaptic boutons were examined and imaged using a JEM-1400 transmission electron microscope (Jeol) at 80 keV.

### Correlative Light Electron Microscopy (CLEM)

Correlative light and electron microscopy (CLEM) was performed as previously described (Soukup et al., 2016). Third instar larvae were dissected in cold HL3 and fixed for 2 h at 4°C (0.5 % glutaraldehyde, 2 % para-formaldehyde in 0.1 M PBS, pH 7.4). After washing in 0.1 M PBS, samples were stained with DAPI (Sigma). Pre-fixed larval fillets were then branded using a Zeiss LSM 780 equipped with a Mai Tai HP DeepSee laser (Spectra-Physics) at 880 nm with 40 % maximal power output. Z-stacks of the ROI where acquired before and after branding with a 25X water immersion lens (NA 0.8). Samples were then post-fixed (4 % para-formaldehyde, 2.5 % glutaraldehyde in 0.1 M phosphate buffer) overnight at 4°C. The lavae were washed with 0,1 M PBS and afterwards with ddH2O. During the rest of the preparation the larvae were washed after every step with ddH2O until the dehydration steps. Branded larvae were then osmicated for 1 h (1 % OsO_4_ and 1.5 % potassium ferrocyanide). Then, the larvae were incubated in a 0.2 % tannic acid for 30 min followed by a second osmication step (1 % OsO_4_ for 30 min) and subsequently put for 20 min in 1 % thiocarbohydrazide. The larvae were osmicated for a third time (1 % OsO_4_ for 30 min) and incubated overnight in 0.5 % uranyl acetate. Samples were then stained with lead aspartate (Walton’s lead aspartate: 20 mM lead nitrate in 30 mM sodium aspartate, pH 5.5) for 30 min at 60°C. After a final washing step the larvae were dehydrated with solutions of increasing ethanol concentration (30 %, 50 %, 70 %, 90 % and twice with 100 %) followed by two 10 min incubation steps with propylene oxide (PO). Larval fillets were then infiltrated with resin agar 100 (Laborimpex), flat embedded in resin agar 100 and placed at 60°C for 48 h.

For 3D-SEM, the flat resin-embedded larval fillets were cropped into 1 mm^2^ pieces with region of interest in the middle and mounted on an aluminium specimen pin (Gatan) using conductive epoxy (Circuit Works). For approaching the region of interest, aluminium pins were places in a Zeiss Sigma VP SEM equipped with Gatan 3View technology. Once the first branding marks were reached and muscle morphology was recognized by correlating with the light microscopy data,70 nm sections were cut on an ultramicrotome (EM UC7, Leica) and collected on slot grids (Van Loenen Instruments). The sections were imaged using a JEM-1400 transmission electron microscope (Jeol) at 80 kV. Branding marks around the NMJ and DAPI signal were used to correlate the confocal images with the TEM micrographs of the NMJ boutons. Overlay images were generated using ImageJ and Gimp.

### FM 1-43

The labeling and quantification of FM1-43 intensities was performed as previously described (Verstreken et al., 2008). Third instar larvae were dissected in fresh Ca^2+^ free HL3, nerves were cut and larvae subsequently incubated for 1 min in HL3 with 4 µM FM 1-43 (Invitrogen), 1.5 mM CaCl_2_ and 90 mM KCl. Multiple steps of washing with HL3 before imaging removed the non-internalized dye. Images of FM 1-43 were captured with an upright widefield microscope (Nikon Eclipse FN1), fitted with 60X (NA 1.0) water dipping lens and stored using NIS elements. Mean boutonic intensities were determined, after background substraction, using ImageJ.

### Electrophysiology

Current clamp experiments to record EJPs were performed as previously described (Kasprowicz et al., 2014; Slabbaert et al., 2016). Third instar larvae were dissected in Ca^2+^ free HL3 which was subsequently replaced with HL3 with 2 mM CaCl_2_ and nerves were cut below the ventral nerve cord. Motor nerves from muscle 6-7, segment A2 or A3 were stimulated at 10 Hz for 400s at least 50 % above the threshold, using a suction electrode. EJP sets were omitted when the recording did not hold its basal membrane potential throughout the 400 s stimulation paradigm. Signals were amplified using the Axoclamp900A amplifier (Molecular Devices), filtered using a 1 kHz Bessel filter and digitized at 10 kHz using a Digidata 1440A (Molecular Devices). Data storage, processing and analysis was done using Clampfit 10.7 (Molecular Devices). EJP amplitudes were quantified for each of the stimuli over the 400 s stimulation duration. The EJP amplitudes were then binned per 300 stimuli with the exception of the first 150 stimuli. The consecutive EJP amplitudes for each binned data point were normalized to the first binned data point of the first 150 stimuli.

### Western blot

Flies collected separately from three independent crosses were decapitated and heads homogenized with a motorized pestel in lysis buffer (25 mM HEPES, 100 mM NaCl, 1 mM CaCl_2_, 1 % Triton, 1X Complete Protease Inhibitor (Sigma)). After incubation on ice for 30 min, samples were spun down at 10000 g for 10 min and supernatant collected and quantified by Bradford assay (BioRad) in a GloMax Multi Detection Plate Reader (Promega). After boiling in 1X Laemmli buffer with 8 % 2-mercapto-ethanol (Sigma), samples were ran on a NuPage 4-12 % Bis-Tris gel (Thermo Fisher Scientific) and transferred on a nitrocellulose membrane (BioRad) subsequently blocked with 10 % BSA in TBS. Primary antibodies were incubated overnight at 4°C in antibody solution (5 % BSA in 0.05 % TBS-T). Fluorescent secondary antibodies were incubated for 1 h at room temperature in antibody solution. After detection with an iBright imaging system (Thermo Fisher Scientific), fluorescent bands were quantified in ImageJ. EndoA fluorescence was normalized to GAPDH fluorescence.

The following antibodies were used: guinea pig anti-EndoA (GP69) [1:5000 (Verstreken et al., 2002)], rabbit anti-GAPDH [1:2000 (Abcam)], mouse anti-ENDOA1 [1:500 (Santa Cruz)], mouse anti-Dynamin [1:1000 (BD Biosciences)], rabbit anti-GST [1:2000 (Life Technologies)]. Alexa Fluor-488/Alexa Fluor-647 conjugated secondary antibodies [1:1000 (Invitrogen)].

### Recombinant protein production and purification

GST-tagged recombinant proteins were produced in competent BL21(DE3) *E.coli* by induction with 1 mM IPTG (Thermo Fisher Scientific) at 37°C for 4 h followed by additional overnight incubation at 25°C. Bacteria were collected by centrifugation at 8000 g, resuspended in lysis buffer (4 mM DTT, 1 mM EDTA, 1X Protease Inhibitor EDTA-free in 1X PBS. pH 6.5-8) and lysed using a high pressure homogenizer operating at a pressure between 15000-20000 bar (Emulsiflex C5, Avestin). Homogenized samples were incubated with 1X DNaseI and spun down at 25000 g for 40 min at 4°C.

Purification was perfomerd using affininty chromatography by loading the supernatant on two 5 ml GSTrap HP column (GE Healthcare) mounted in series and operated at 5 ml/min with an ÄKTA Pure 25 system (GE Healthcare). After binding, the column was washed with 10 column volumes of PBS followed by 5 column volumes of PreScission buffer (20 mM Tris, 100 mM NaCl, 1 mM EDTA, 1 mM DTT, pH 8). Cleavage of the GST tag was perfomed on column at room temperature for 4 h with 500 U of PreScission Protease (GE Healthcare) diluted in PreScission buffer. Cleaved protein was collected in PBS by peak fractionation, while the remaining uncleaved protein was eluted with 5 colomn volumes of 100 % elution buffer (20 mM Tris, 10 mM Gluthatione reduced, pH 8).

Input material and collected fractions were analysed by SDS-PAGE on NuPage 4-12 % Bis-Tris gel (Thermo Fisher Scientific) and stained with Coomassie.

EndoA protein concentrations were estimated by absorbance using the calculated molar extinction coefficient of 17670 M^-1^cm^-1^ for the *Drosophila* proteins, and 17545 M^-1^cm^-1^ for the human proteins.

### Biphysical/structural characterization of recombinant proteins

Prior to biophysical/structural characterization experiments, purified proteins were subjected to gel filtration to isolate the peak corresponding to dimeric EndoA. A Superdex 200 increase 30/100 column (GE Healthcare) was operated at a flow rate of 0.5 ml/min and buffered with 20 mM MOPS, 150 mM NaCl pH 7. The elution volume of the dimer peak was previously determined by SEC-MALS.

Thermal stability was determined through thermal denaturation (from 20°C to 95°C at an increment rate of 0.3°C/min) monitored by fluorescent emission intensity upon excitation at 266 nm. Measurements were obtained in an Uncle instrument (Unchained Labs). EndoA^WT^ was concentrated to ∼3 mg/ml (Amicon Ultra centrifugal filters) and filtered through a 0.22 μm filter (Millex). Thermal stability was tested in the presence of 1 mM EDTA or 1 mM CaCl_2_. Measurements were done in triplicate and average thermograms were displayed by plotting the barycentric mean (BCM) and Static light scattering (SLS) at 266 nm.

Fourier Transform InfraRed (FTIR) spectroscopy was performed on a Invenio spectrometer (Bruker) equipped with a BioATR-II measuring cell (Bruker). Proteins were concentrated to ∼2 mg/ml and filtered through a 0.22 μm filter (Millex). Recordings were performed at 25°C. The temperature was controlled by a thermostatic water bath. Spectra were acquired with a resolution of 4 cm^-1^ and with a total of 120 scans per spectra averaged to improved the signal-to-noise ratio.

Dynamic light scattering (DLS) measurements were performed in a DynaPro DLS plate reader (Wyatt). Proteins were concentrated to ∼2 mg/ml, filtered through a 0.22 μm filter (Millex) and loaded on a low binding 394-well black plate with transparent bottom (Costar). The autocorrelation of scattered light intensity was averaged over 5 recordings to obtain single data points. Wyatt Dynamics software (v7) was used to calculate the hydrodynamic radius.

For Small angle X-ray Scattering (SAXS) purified *Drosophila* EndoA and human ENDOA1 proteins were run though a Superdex200 10/300 increase chromatographgy column (GE Healthcare) to isolate the peak corresponding to the dimeric protein. The column was buffered in 20 mM MOPS, 150 mM NaCl, 1 mM DTT, pH 7 or 20 mM MOPS, 150 mM NaCl, 1 mM DTT, 1 mM CaCl_2_, pH 7. Dimeric proteins were concentrated with an Amicon Ultra centrifugal filter to a concentration of at least 6 mg/ml. Concentrations were determined by absorbance. SAXS data were collected according to Supplemental Table 1. The sample of *Drosophila* EndoA^D265A^ mutant was measured at the BM29 beamline of the ESRF synchrotron (Grenoble, France) (Pernot et al., 2013), while all other samples were measured at the SWING beamline of the SOLEIL synchrotron (Paris, France) (Thureau et al., 2021). Measurements were performed using the HPLC-SEC setup available at the beamlines by injecting 30-50 µl of 6-11 mg/ml of protein onto an Advanced BioSEC 300Å 2.7/300 column. The column was pre-equilibrated with a buffer containing 20 mM MOPS (pH 7), 150 mM NaCl, 1 mM DTT, 5 % glycerol and either no calcium or 1 mM CaCl_2_. The flow rate was set at 0.3 ml/min for the measurement performed at the BM29 beamline and at 0.35 ml/min for the ones measured at the SWING beamline. The radial averaging was performed using BsxCuBE (Pernot et al., 2013) for the data collected at the ESRF synchrotron and using FOXTROT (Perez J., Bizien T., 2022) for those collected at the SOLEIL synchrotron. Buffer subtractions were performed using CHROMIXS (Panjkovich and Svergun, 2018) and the averaged data, corresponding to the peak of interest, were further processed using the ATSAS software package (Petoukhov et al., 2012). The molecular weight estimations were taken from the Bayesian assessment method (Hajizadeh et al., 2018), while the Kratky plots were generated using the ATSAS program PRIMUS (Konarev et al., 2003)).

### Co-immunoprecipitation

50 μl of slurry Dynabeads Protein G (Thermo Fisher) were coupled with 5 μg of mouse monoclonal anti-Dynamin antibody (BD Biosciences) rotating for 1 h at room temperature. Beads were washed twice with wash buffer (50 mM Tris, 150 mM NaCl, pH 7.5) and incubated overnight at 4°C with 1 mg of total *Drosophila* (*w^1118^*) head lysate diluted in wash buffer to a final volume of 400 μl. After washing the beads twice with wash buffer, they were incubated for 4 h at room temperature with 30 μg of purified GST-EndoA^WT^, GST-EndoA^D265A^ or GST-EndoA^D265R^. Following washes, the beads were resuspended in wash buffer and loading dye (4X Laemlli buffer (Bio-Rad) + 2-mercapto ethanol (Sigma)), boiled for 10 min at 95°C and loaded on NuPage 4-12 % Bis-Tris gel (Thermo Fisher Scientific). After transferring the proteins on a nitrocellulose membrane (BioRad), the membrane was processed for Western blotting.

Control Co-IP were performed by incubating 50 μl of slurry Dynabeads Protein G, not coupled with anti-Dynamin antibody, with 1 mg of total head lysate followed by incubation with 30 μg of purified GST-EndoA^WT^. Incubation times were the same as for the other conditions.

GST-EndoA proteins used for this assay were purified as described in the ‘recombinant proteins production and purification’ paragraph, with the exception that the GST tag was not cleaved. Proteins were eluted from the 5ml GSTrap HP column with elution buffer (20 mM Tris, 10 mM Gluthatione reduced, pH 8) and dialyzed overnight at 4°C in 50 mM Tris, 150 mM NaCl, pH 7.5. GST-EndoA proteins concentrations were estimated by absorbance using the calculated molar extinction coefficient of 60780 M^-1^cm^-1^.

### Electroretinograms (ERGs)

ERGs were recorded from 1 to 3-day-old flies as previously described (Soukup et al., 2016). Flies were immobilized on glass microscope slides, by use of double-sided tape. For recordings, glass electrodes (borosilicate, 1.5 mm outer diameter) filled with 3 M NaCl were placed in the thorax as a reference and on the fly eye for recordings. Responses to repetitive light stimuli were recorded using Axosope 10.7 and analyzed using Clampfit 10.7 software (Molecular Devices) and Igor Pro 6.37.

### Light induced neurodegeneration and histology

Light induced retinal degeneration induced by placing 1 to 3-day-old flies under continuous illumination (1300 lux) (Soukup et al., 2013). Batches of flies were also kept in darkness for 7 days at 25°C. Flies were used for ERG data acquisition, or processed for histological staining of the retina.

Histological sections of the retina were prepared by decapitating heads and fixing them in 4 % para-formaldehyde and 2.5 % glutaraldehyde in 0.1 M PBS overnight at 4°C or until further processing. Heads were then osmicated in 2 % OsO_4_ for 2 h and subsequently incubated in 4 % uranyl acetate for 1 h. After dehydration using an ethanol series, heads were embedded in hard resin (Agar 100, Laborimpex) and semi-thin (1.5 μm) sections were cut on a microtome (EM UC7, Leica) and stained on a heating block with a 1 % toluidine blue (Merck) solution including 2 % Borax for 90 s at 60°C. The stained sections were mounted with Eukit Quick-hardening mounting medium (Sigma). Histological sections were analyzed using the Leica DM2500 M microscope equipped with a 40X lens.

### Gene editing of iPSC

*SH3GL2* p.G276V was engineered in the “Ctrl65” iPSC line (SFC065, (Baumann et al., 2021)), by means of CRISPR/Cas9 gene editing using a split puromycin recombination reporter (Flemr and Bühler, 2015). One million cells were seeded per well of a 6-well plate in the presence of Rho kinase inhibitor the day before transfection (Miltenyi Biotech; 130-106-538). The next day, the cells were transfected using Lipofectamine Stem (Life Technologies; STEM00015) with a mix of 1 µg donor single stranded oligodeoxynucleotide, 0.5 µg pRR-Puro plasmid (Addgene plasmid #65853) specific for the guide RNA (gRNA) spacer sequence and 1 µg px458 encoding for a guide RNA targeting the mutation site (Addgene plasmid #65853; (Ran et al., 2013)). Two days post-transfection, cells were treated with 0.5 µg/ml puromycin for 48 h in the presence of Rho kinase inhibitor. Ten to fourteen days after transfection, resistant colonies were manually picked, expanded, and molecularly characterized by means of PCR, Sanger sequencing and digital droplet PCR to discard large deletions. Clones showing the desired genotype were karyotyped by means of comparative genomic hybridization (CGH) and stained for pluripotency markers.

### iPSC differentiation

On day -1, 300.000 hiPSC/cm^2^ were seeded on Matrigel coated 6-well plate wells in mTeSR-Plus medium supplemented with 10 µM Rho kinase inhibitor (RI). On day 0, medium was switched to knockout serum replacement medium (KSR) containing DMEM/F-12, 15 % knockout serum replacement, GlutaMAX, Penstrep, non-essential amino acids and 10 μM β-mercaptoethanol (all from Life Technologies) supplemented with LDN193189 (500 nM, Sigma), SB431542 (10 μM, Tocris), SHH-C24II (100 ng/ml, Miltenyi Biotec), Purmorphamine (2 μM, Sigma). CHIR99021 was added to the medium from day 3 to day 13 (CHIR; 3 μM, Stemcell Technologies). From day 4, KSR was gradually shifted to Neurobasal/0.5X B27 supplement without vitamin A, 0.5X N2, GlutaMAX, Penstrep, non-essential amino acids at the rate of 1/3 every 2 days for 4 days, and to ¼ for 3 days. SB431542, SHH-C24II, and Purmorphamine were withdrawn from the medium at day 7. FGF8b (100 ng/mL, R&D) was added to the medium from day 9 until day 16. At day 18, cells were switched to terminal differentiation medium consisting of Neurobasal-A/1xB27 supplement without vitamin A, GlutaMAX, Penstrep containing BDNF (brain-derived neurotrophic factor, 10 ng/ml; R&D), ascorbic acid (0.2 mM, Sigma), GDNF (glial cell line-derived neurotrophic factor, 10 ng/ml; R&D), dibutyryl cAMP (0.2 mM; Sigma), SR11237 (100 nM, Tocris), and DAPT (10 μM; Tocris). On day 20, ventral midbrain neural progenitors were cryopreserved and quality controlled. Neural progenitor cells were terminally differentiated on coverslips previously coated with poly-D-lysine (50 µg/mL)/mouse laminin (1 µg/mL) in terminal differentiation medium for additional 35 days.

### Analysis of LC3B puncta in TH^+^ neurites

Terminally differentiated vmDAn (55-60 days) were incubated for 15 min in physiological solution (20 mM HEPES pH 7.4, 140 mM NaCl, 4.7 mM KCl, 2.5 mM CaCl_2_, 1.2 mM MgSO_4_, 1.2 mM KH_2_PO_4_, 11 mM glucose) and subsequently fixed with 4 % para-formaldehyde, processed for immunostaining and imaged. Z-stack confocal images were analyzed with ImageJ in a semi-automated way. The ‘Skeletonize’ plugin was ran on the TH channel and the total length of TH^+^ neurites was measured. The skeleton was converted to a mask within which LC3B puncta were counted using the ‘Analyze particle’ plugin. The number of LC3B puncta within the mask were normalized by the total length of TH^+^ neurites and plotted as LC3B puncta/TH^+^ unit length.

### Statistics

GraphPad Prism 9.3 (San Diego, USA) was used to determine statistical significance. Datasets were tested for normal distribution using the D’Agostino-Person Omnibus and Shapiro-Wilk normality tests. Normally distributed data were tested with parametric tests: when two datasets were compared, the Student’s *t*-test was used, and when more than two datasets were compared, a one-way analysis of variance test (ANOVA) followed by a *post hoc* Tukey test was used. For non-normally distributed datasets, Mann-Whitney test was used when comparing two datasets, and an ANOVA Kruskal-Wallis test followed by a Dunn’s *post hoc* test was used for multiple datasets. When multiple parameters were compared (genotypes and treatments) a two-way ANOVA was used, followed by a *post hoc* Tukey test or Šidàk test for multiple comparison correction. Significance levels are defined as \*\*\**P* < 0.0001, \*\**P* < 0.01, \**P* < 0.05 and ns, not significant. ‘*n*’ in the legends indicates the number of animals used and analyzed. For sptPALM images 2 separate NMJs were imaged per animal, while for the confocal imaging and single molecule localization studies, 4 different NMJs were imaged in each animal. Data are plotted as mean ± SEM.

---

**Supplemental Table 1:**
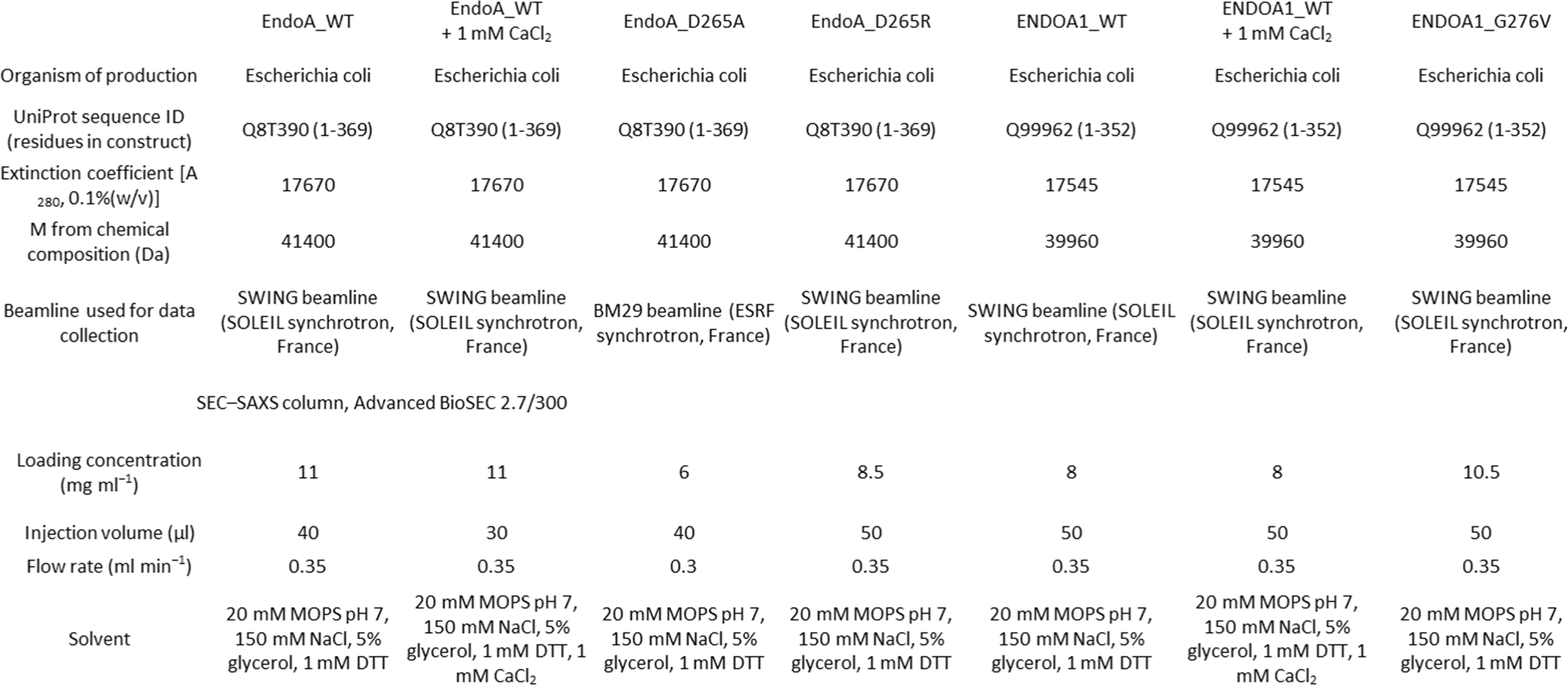
Experimental conditions of SAXS data collection.

## Acknowledgements

We thank VIB-KU Leuven Center for Brain and Disease Research light microscopy unit for their help. We thank the members of the Verstreken lab for helpful discussions and comments. Dr Tristan Wallis (The University of Queensland, Australia) provided edited DBSCAN code for single molecule localization cluster analysis. Dr Sam Duwe (University of Hasselt, Belgium) provided technical expertise for the acquisition of single molecule imaging on the ELYRA. A Zeiss LSM 780 – SP Mai Tai HP DS (Cell Tissue Imaging Cluster (CIC)) used for confocal images for CLEM is supported by Hercules AKUL/11/37 and FWO G.0929.15 to Pieter Vanden Berghe, University of Leuven. Research support was provided by ERC consolidator grants (to P.V.), the Chan Zuckerberg Initiative (to P.V.), FWO Vlaanderen (to P.V, W.V., F.R., J.S., E.M., C.C., N.L.), Initiative d’Excellence de l’Université de Bordeaux (IDEX Neurocampus Chair, to S.F.S.), GPR BRAIN_2030 (to S.F.S.) and the Region Nouvelle-Aquitaine (to S.F.S.). The Switch laboratory received funding from the Flanders Institute for Biotechnology (VIB) and KU Leuven. This research was also funded in part by Aligning Science Across Parkinson’s [ASAP-000430] through the Michael J. Fox Foundation for Parkinson’s Research (MJFF). A.T.B. is supported by the EMBO long-term postdoctoral fellowship (ALTF-1034-2018) and Race against dementia – Dementia Australia postdoctoral fellowship. P.V. is an alumnus of the FENS-Kavli Network of Excellence.

## Author Contributions

Conceptualization, A.T.B., M.D., S.K., S.F.S. and P.V. Methodology, A.T.B., M.D., S.K., and P.V. Investigation: A.T.B., M.D., S.K., C.C., J.S., S.F.G., N.S., N.L., E.M., S.K., K.V., and N.G. Formal Analysis: A.T.B., M.D., S.K., J.S., J-B.S., F.A.M., A.E., W.V., F.R., and J.S. Writing, A.T.B., M.D., S.K., and P.V. Funding Acquisition, A.T.B., C.C., N.L., E.M., W.V., F.R., J.S. and P.V. Supervision, P.V.. All co-authors read and edited the manuscript. The co-first authors are listed in alphabetical order and they explicitly state the equal nature of their contribution and thus, the interchangeability of the order they appear on this paper.

