## Supplemental Figures and legends for "EndophilinA-dependent coupling between activity-dependent calcium influx and synaptic autophagy is disrupted by a Parkinson-risk mutation"

Figure S1

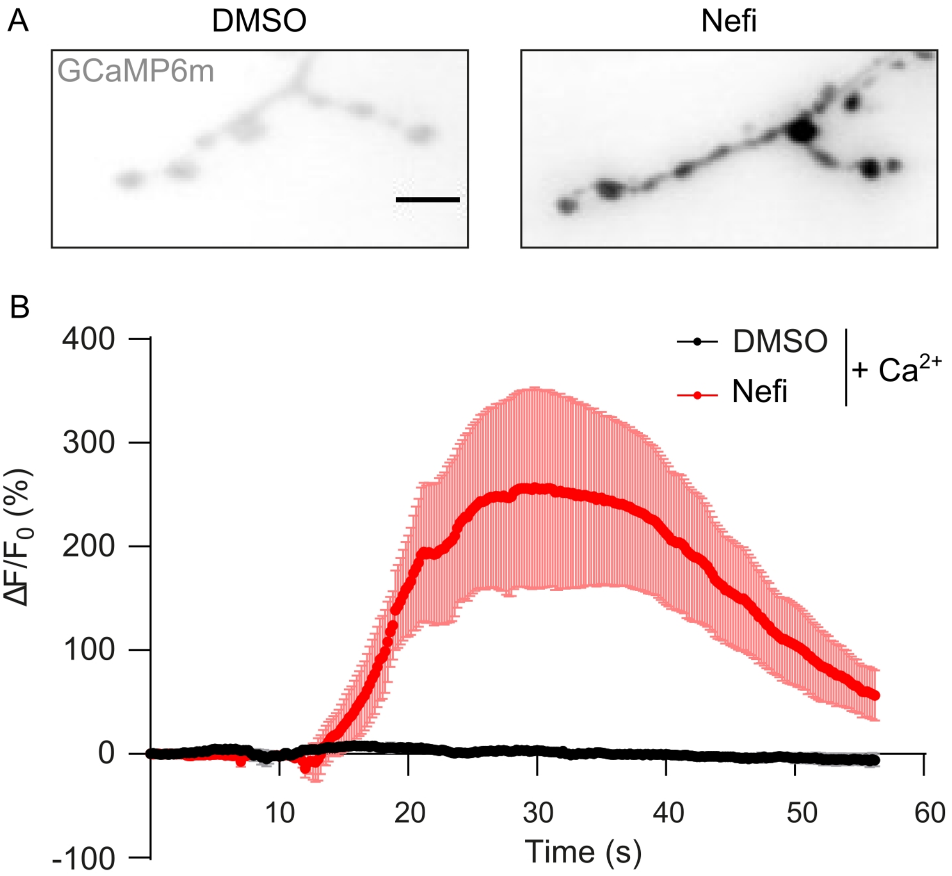
A
DMSO
Nefi
GCaMP6m
B
400
300
200
100
0
-100
ΔF/F0 (%)
10
20
30
40
50
60
Time (s)
DMSO
Nefi
+ Ca2+

Figure S2

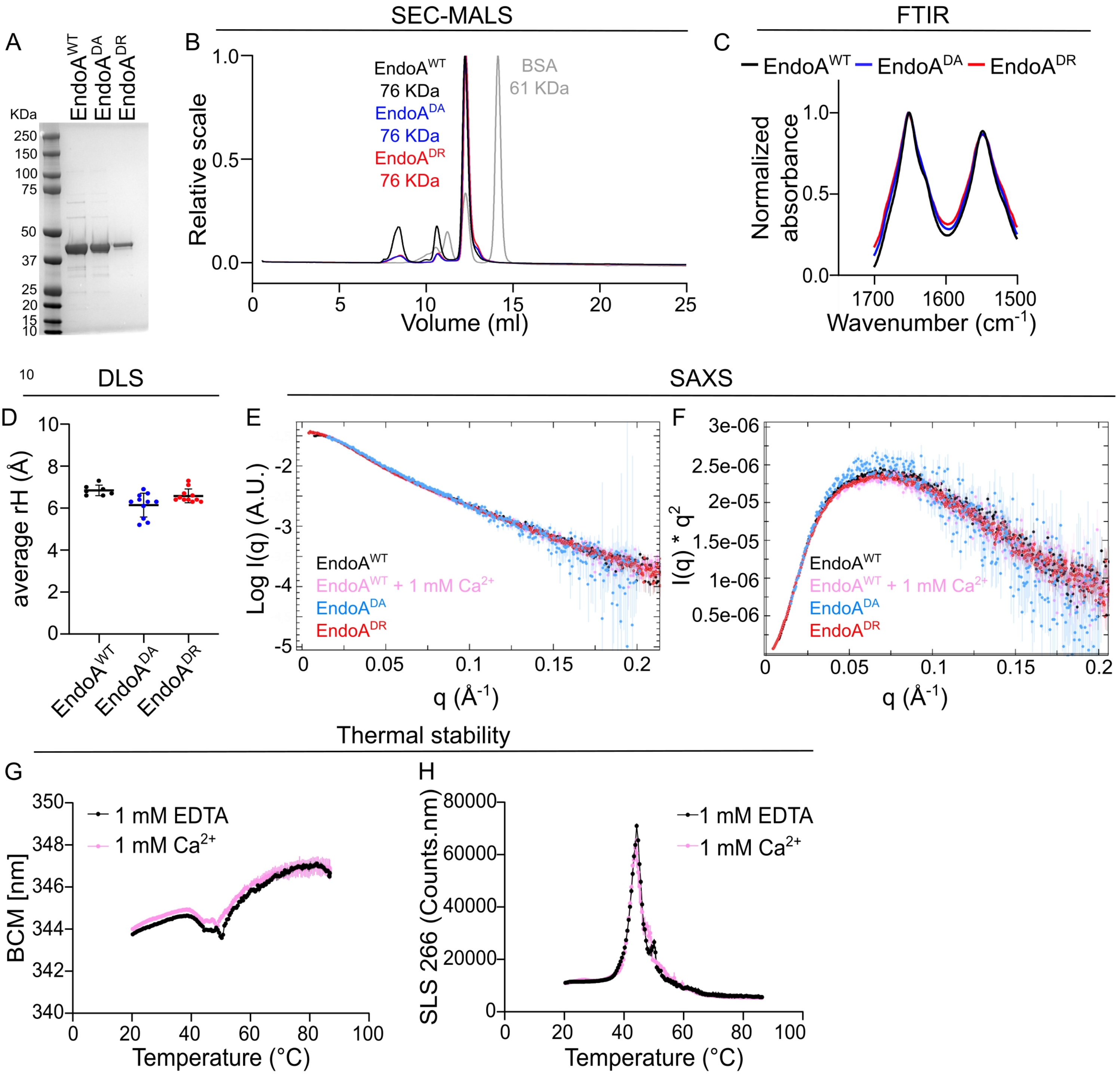

SEC-MALS
FTIR
A
B
C
EndoA$^{WT}$
EndoA$^{DA}$
EndoA$^{DR}$
KDa
250
150
100
75
50
37
25
20
15
10
Relative scale
EndoA$^{WT}$ 76 KDa
EndoA$^{DA}$ 76 KDa
EndoA$^{DR}$ 76 KDa
BSA 61 KDa
Volume (ml)
Normalized absorbance
Wavenumber (cm$^{-1}$)
10
DLS
SAXS
D
E
F
average rH (Å)
Log I(q) (A.U.)
I(q) * q$^2$
EndoA$^{WT}$ + 1 mM Ca$^{2+}$
q (Å$^{-1}$)
Thermal stability
G
H
BCM [nm]
SLS 266 (Counts.nm)
1 mM EDTA
1 mM Ca$^{2+}$
Temperature (°C)

Figure S3

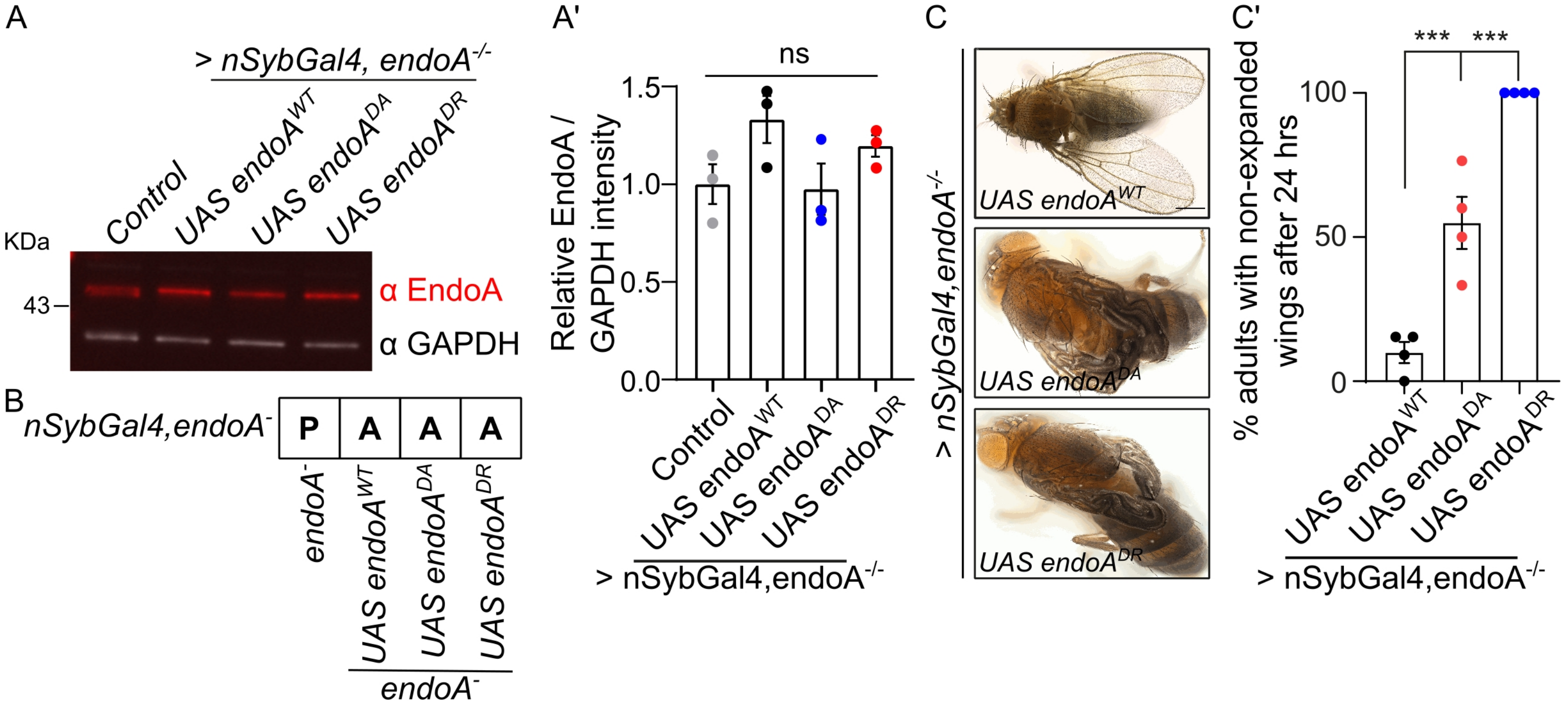
A
> nSybGal4, endoA-/-
Control
UAS endoAWT
UAS endoADA
UAS endoADR
KDa
43
α EndoA
α GAPDH
A'
ns
Relative EndoA / GAPDH intensity
1.5
1.0
0.5
0.0
Control
UAS endoAWT
UAS endoADA
UAS endoADR
> nSybGal4,endoA-/-
B
nSybGal4,endoA-
P
A
A
A
endoA-
UAS endoAWT
UAS endoADA
UAS endoADR
endoA-
C
> nSybGal4,endoA-/-
UAS endoAWT
UAS endoADA
UAS endoADR
C'
***
***
% adults with non-expanded wings after 24 hrs
100
50
0
UAS endoAWT
UAS endoADA
UAS endoADR
> nSybGal4,endoA-/-

Figure S4

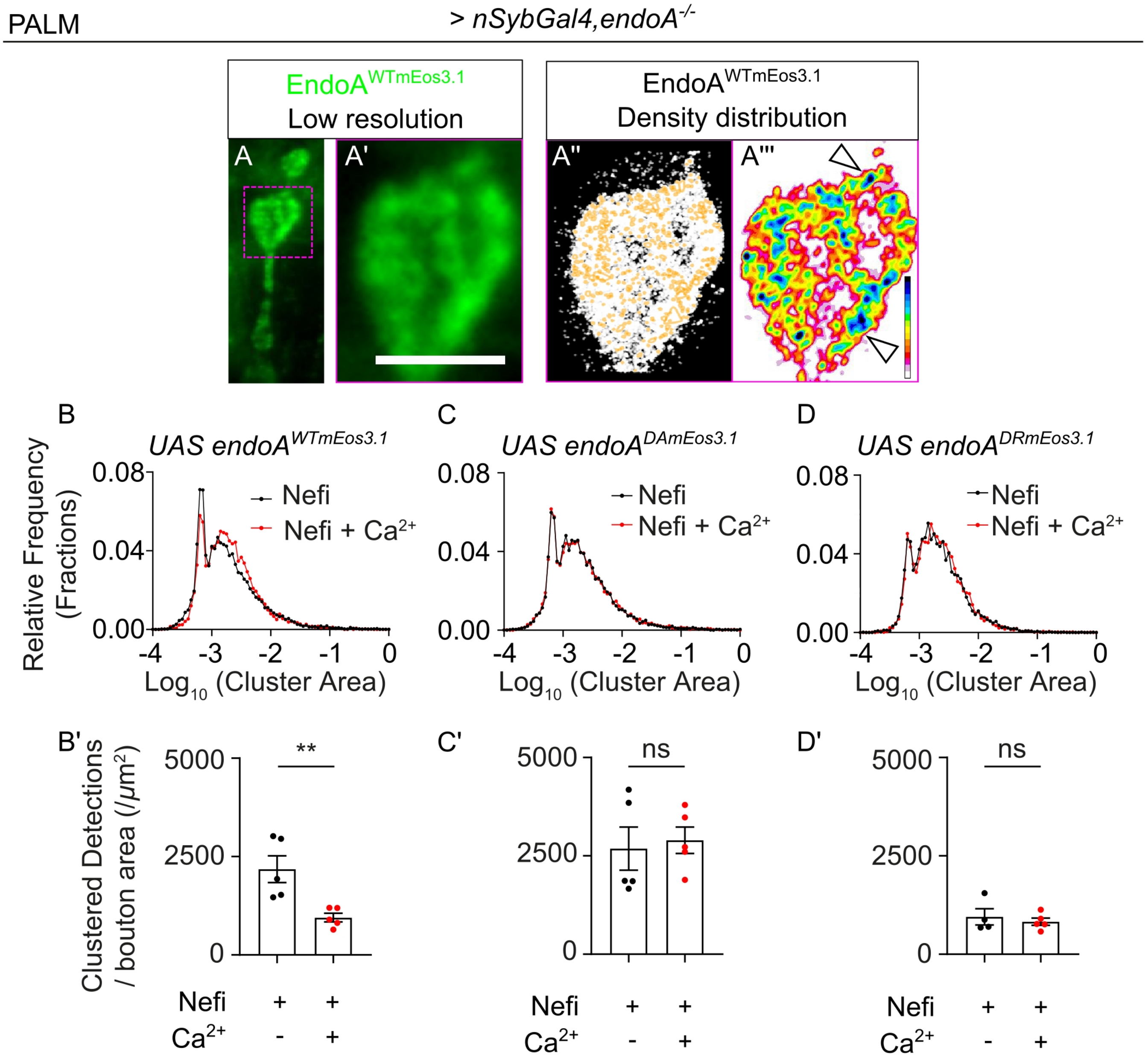

PALM
> nSybGal4,endoA-/-
EndoA WTmEos3.1
Low resolution
EndoA WTmEos3.1
Density distribution
A
A'
A''
A'''
B
UAS endoA WTmEos3.1
C
UAS endoA DAmEos3.1
D
UAS endoA DRmEos3.1
Relative Frequency (Fractions)
Nefi
Nefi + Ca2+
0.08
0.04
0.00
-4 -3 -2 -1 0
Log10 (Cluster Area)
B'
C'
D'
Clustered Detections / bouton area (/µm2)
5000
2500
0
**
ns
ns
Nefi + +
Ca2+ - +

Figure S5

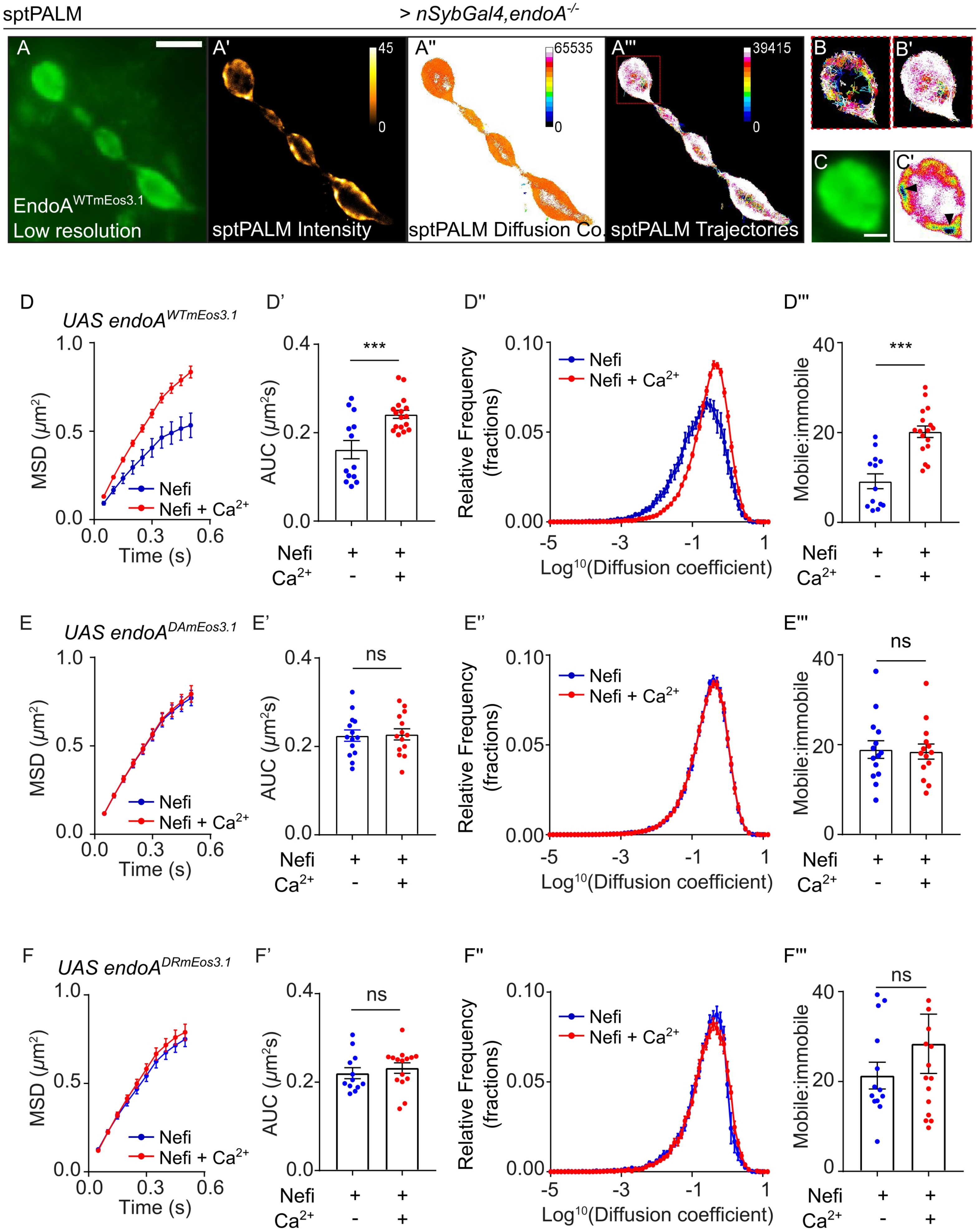

sptPALM
> nSybGal4,endoA-/-
A
A'
A''
A'''
B
B'
C
C'
EndoA WTmEos3.1
Low resolution
sptPALM Intensity
sptPALM Diffusion Co.
sptPALM Trajectories
45
0
65535
0
39415
0
D
UAS endoA WTmEos3.1
D'
D''
D'''
E
UAS endoA DAmEos3.1
E'
E''
E'''
F
UAS endoA DRmEos3.1
F'
F''
F'''
MSD (μm²)
Time (s)
Nefi
Nefi + Ca2+
AUC (μm²s)
Relative Frequency (fractions)
Log10(Diffusion coefficient)
Mobile:immobile
Ca2+
***
ns

Figure S6

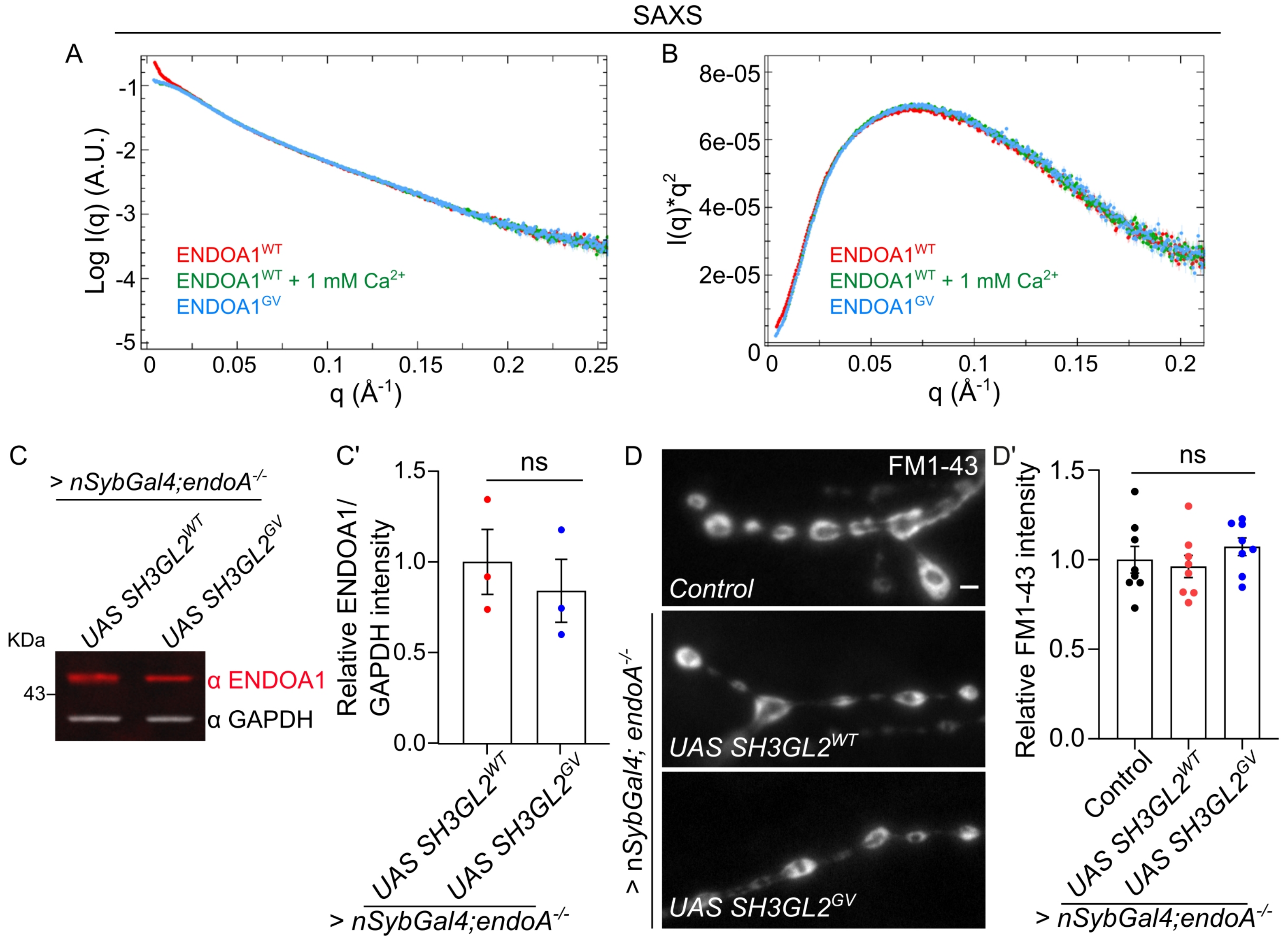

SAXS
A
Log I(q) (A.U.)
q (Å-1)
ENDOA1WT
ENDOA1WT + 1 mM Ca2+
ENDOA1GV
B
I(q)*q2
q (Å-1)
ENDOA1WT
ENDOA1WT + 1 mM Ca2+
ENDOA1GV
C
> nSybGal4;endoA-/-
UAS SH3GL2WT
UAS SH3GL2GV
KDa
43
α ENDOA1
α GAPDH
C'
Relative ENDOA1/ GAPDH intensity
ns
UAS SH3GL2WT
UAS SH3GL2GV
> nSybGal4;endoA-/-
D
FM1-43
Control
UAS SH3GL2WT
UAS SH3GL2GV
> nSybGal4; endoA-/-
D'
Relative FM1-43 intensity
ns
Control
UAS SH3GL2WT
UAS SH3GL2GV
> nSybGal4;endoA-/-

Figure S7

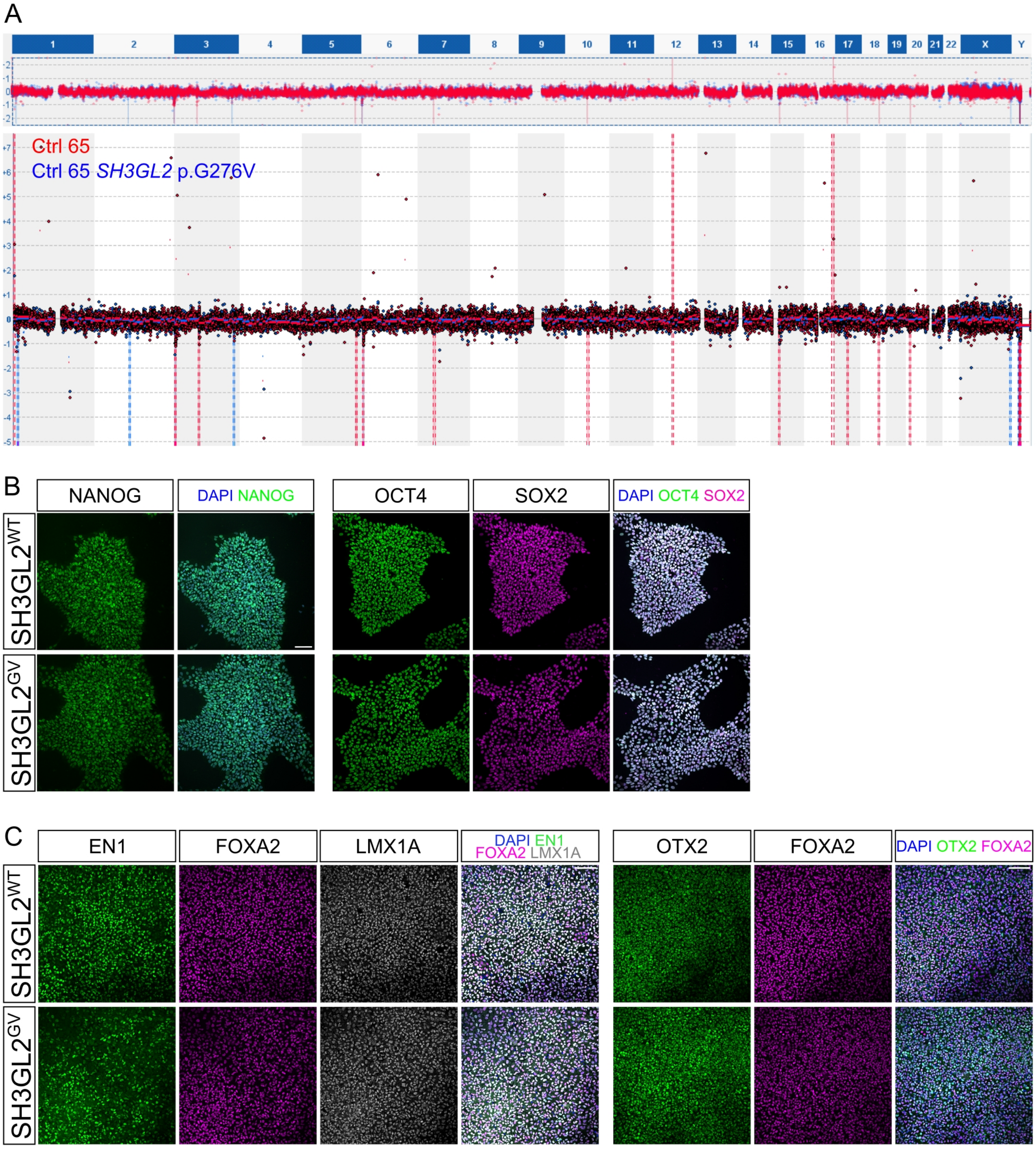
A
Ctrl 65
Ctrl 65 SH3GL2 p.G276V
B
NANOG
DAPI NANOG
OCT4
SOX2
DAPI OCT4 SOX2
SH3GL2WT
SH3GL2GV
C
EN1
FOXA2
LMX1A
DAPI EN1 FOXA2 LMX1A
OTX2
FOXA2
DAPI OTX2 FOXA2
SH3GL2WT
SH3GL2GV

### Supplementary Figure Legends

**Supplementary Figure 1 $Ca^{2+}$ channel agonist, Nefiracetam increases $Ca^{2+}$ influx in synaptic boutons, related to Figure 1**

(A) Representative images of NMJs expressing GCaMP6m under a pan-neuronal driver (*nSyb-Gal4*). Animals were perfused in either HL3 solution supplemented with DMSO, NAS (100 µM) and $CaCl_2$ (2 mM) or in HL3, Nefiracetam (Nefi) (150 µM), NAS (100 µM) and $CaCl_2$ (2 mM).

(B) Quantification of GCaMP6m fluorescence upon Nefiracetam-induced calcium influx. Number of animals ≥ 3 per condition.

**Supplementary Figure 2 EndoA mutants do not show large conformational changes, related to Figure 2**

We wondered if the differential association of the EndoA mutants with Dynamin could be explained by conformational differences, as speculated in literature (Chen et al., 2003; Zhang et al., 2012a).

(A) Coomassie staining of SDS-PAGE gel to visualize purified *Drosophila* $EndoA^{WT}$, $EndoA^{D265A}$ and $EndoA^{D265R}$ after size exclusion chromatography (SEC). Expected size of monomeric EndoA proteins: ~41 KDa.

(B) Size Exclusion Chromatography coupled with Multiple Angle Light Scattering (SEC-MALS) chromatograms of purified *Drosophila* $EndoA^{WT}$, $EndoA^{D265A}$ and $EndoA^{D265R}$ ran on a Superdex 200 increase 30/100 column show that the purified proteins are homogeneous and reveal that D265 mutations do not affect the ability of EndoA to dimerize. EndoA dimer peak elutes at ~12 ml. Expected size of dimeric EndoA: ~ 82 KDa. Experimentally measured seizes reported on graph. EndoA proteins chromatograms were normalized against the monomeric peak of BSA.

(C) Normalized absorbance specta of purified *Drosophila* $EndoA^{WT}$, $EndoA^{D265A}$ and $EndoA^{D265R}$ measured by Fourier Transform Infrared (FTIR) spectroscopy show no differences in secondary structure composition between mutant and wild type proteins. Wavenumbers between 1700-1600 $cm^{-1}$ correspond to the amide I peak, while the region between 1600-1500 $cm^{-1}$ corresponds to the amide II peak.

(D) Average hydrodynamic radii of *Drosophila* $EndoA^{WT}$, $EndoA^{D265A}$ and $EndoA^{D265R}$ measured by Dynamic Light Scattering (DLS) indicate no large differences between the tested proteins. Error bars represent mean ± SEM. ≥ 7 measurements per protein from two independent experiments.

(E-F) To further explore the possibility of a conformational change, we performed Small Angle X-ray Scattering (SAXS) measurements. (E) Scattering profiles obtained from SAXS measurements of *Drosophila* $EndoA^{WT}$, $EndoA^{D265A}$ and $EndoA^{D265R}$ are fully superimposable, indicating that no large conformational changes can be observed between $EndoA^{WT}$, $EndoA^{D265A}$ and $EndoA^{D265R}$.

(F) Kratky plots of *Drosophila* $EndoA^{WT}$, $EndoA^{D265A}$ and $EndoA^{D265R}$. Bell-shaped curves indicate that the analyzed proteins are compact and well folded.

We also wondered if the presence of $Ca^{2+}$ would induce conformational changes. The scattering profile of EndoA$^{WT}$ in 20 mM MOPS +150 mM NaCl +1 mM DTT +5 % glycerol in the presence of 1 mM $CaCl_2$ is similar to that of the EndoA$^{WT}$ in the absence of 1 mM $CaCl_2$ (E). In addition, thermal stability (G-H) of EndoA$^{WT}$ with 1 mM $CaCl_2$ or 1 mM EDTA are similar.
(G) Thermal stability is displayed as Barycentric mean (BCM) of the instrinsic fluorescence signal of *Drosophila* EndoA$^{WT}$ plotted over a thermal ramp (20-95°C). Lines represents average of three measurements. Melting temperatures ($T_m$) are calculated from the first derivarive of the BCM. $T_m$ for EndoA$^{WT}$ in 1 mM EDTA and 1 mM $CaCl_2$ are respectively 51°C and 49.9°C.
(H) Aggregation propensity is shown by Static light scattering (SLS) measured at 266 nm of *Drosophila* EndoA$^{WT}$ plotted over a thermal ramp (20-95°C). Lines represents average of three measurements.
These results suggest that both mutating D265 or $Ca^{2+}$ do not cause large conformational changes.

**Supplemental Figure 3 Expression of mutant EndoA gives rise to weak adult flies, related to Figure 2**

(A) Westen blot of control (*nSyb-Gal4/+*) and *endoA*$^{-/-}$ mutants expressing *endoA*$^{WT}$, *endoA*$^{D265A}$ and *endoA*$^{D265R}$ under a pan-neuronal driver (*nSyb-Gal4*). Blots were probed with anti-EndoA and anti-GAPDH antibodies. (A') Quantification of EndoA signal intensity relative to GAPDH intensity. Error bars represent mean ± SEM; statistical significance calculated with an ordinary one-way ANOVA with Tukey's multiple comparison test: ns, not significant. Experiment performed in three independent biological replicates.
(B) Table indicating neuronal expression (*nSyb-Gal4*) of *endoA*$^{WT}$, *endoA*$^{D265A}$ and *endoA*$^{D265R}$ in *endoA*$^{-/-}$ mutants. 'A' and 'P' indicate survival to 'adult stage' and 'pupa stage', respectively.
(C) Representative images of adult *endoA*$^{-/-}$ *Drosophila* expressing *endoA*$^{WT}$, *endoA*$^{D265A}$, and *endoA*$^{D265R}$ under the control of the pan-neuronal driver *nSyb-Gal4*. *endoA*$^{D265A}$ and *endoA*$^{D265R}$ adult flies have marked phenotype of non-expanded wings, days after eclosion. Scale bar: 1 mm. (C') Quantification of percentage of adult *Drosophila* with non-expanded wings 24 hours after eclosion across the three genotypes. Statistical significance calculated with an ordinary one-way ANOVA with Tukey's multiple comparison test: *** $P < 0.001$, $n \geq 51$ animals per genotype across 4 independent experiments.

**Supplementary Figure 4 Nanoscale localization of EndoA mutants is unchanged by synaptic activity, related to figure 4**

(A-A''') Transgenic *endoA*$^{-/-}$ larvae expressing *endoA*$^{WT\text{::mEos3.1}}$, *endoA*$^{D265A\text{::mEos3.1}}$ or *endoA*$^{D265R\text{::mEos3.1}}$ (under the pan-neuronal driver *nSyb-Gal4*) were imaged using single molecule localization photoactivated localization microscopy (PALM) at 20 Hz. Low resolution images of NMJ (A) and zoomed in individual bouton (A') expressing *endoA*$^{WT\text{::mEos3.1}}$. Scale bar: 2 µm. (A'') Single molecule localization clusters of the same zoomed in bouton obtained post-processing from photo-converted movie. (A''') Cluster map colour-coded for cluster size and density distribution of EndoA$^{\text{::mEos3.1}}$ generated by density-

based spatial clustering of applications with noise (DBSCAN) analysis. Arrowhead indicate EndoA nanodomains. Fluorescence intensity shown using indicated scale (0-65535).
(B-D) Relative frequency distribution of cluster area of of $endoA^{WT\text{::mEos3.1}}$, $endoA^{D265A\text{::mEos3.1}}$ and $endoA^{D265R\text{::mEos3.1}}$ in non-stimulated (30 min incubation in HL3, Nefi (10 µM) and NAS (100 µM) solution and stimulated conditions (30 min incubation in HL3 solution containing Nefi (10 µM), NAS (100 µM) and $CaCl_2$ (1 mM)). Error bars represent mean ± SEM; n ≥ 5 larvae (10 NMJs) per genotype.
(B'-D') Quantification of average clustered detections per unit bouton area in the different genotypes in non-stimulated and stimulated conditions. Error bars represent mean ± SEM; statistical significance was calculated with an student *t*-test two-tailed unpaired distribution: ** $P < 0.01$, ns, not significant, n ≥ 5 larvae (20 NMJs) per genotype.

**Supplementary Figure 5 $Ca^{2+}$ influx changes the mobility of EndoA, related to Figure 4**
(A-A''') Representative low resolution image of transgenic $endoA^{-/-}$ larvae expressing $endoA^{\text{::mEos3.1}}$ (under the pan-neuronal driver *nSyb-Gal4*) (A), super-resolved average intensity map (A'), super-resolved diffusion coefficient map (A'') and super-resolved trajectories map (A'''). Fluorescence intensities shown using indicated scale in A' and A''. (B, B') Show trajectories of single EndoA molecules classified as immobile and mobile population.
(C-C') Low resolution image of a zoomed in bouton (C) and sptPALM super-resolved average intensity map (C') diplaying clear accumulation of $\text{EndoA}^{\text{::mEos3.1}}$ in nanodomains (arrowheads). Scale bar: 2 µm.
(D-F) Change in MSD (µm$^2$) (D, E, F), area under MSD curves (µm$^2$s) (D', E', F'), relative frequency distribution of diffusion coefficients (D'', E'', F'') and ratio of mobile to immobile population (D''', E''', F''') in $endoA^{-/-}$ larvae expressing $endoA^{WT\text{::mEos3.1}}$, $endoA^{D265A\text{::mEos3.1}}$ and $endoA^{D265R\text{::mEos3.1}}$ (under control of *nSyb-Gal4*) in non-stimulated and stimulated conditions. The increase in mobility of EndoA in $endoA^{WT\text{::mEos3.1}}$ due to $Ca^{2+}$ is absent in $endoA^{D265A\text{::mEos3.1}}$ and $endoA^{D265R\text{::mEos3.1}}$. Error bars represent mean ± SEM; statistical significance was calculated with an student *t*-test two-tailed unpaired distribution: *** $P < 0.001$, ns, not significant, n ≥ 5 larvae (20 NMJs) per genotype.

**Supplementary Figure 6 *SH3GL2* coding variant does not affect ENDOA1 conformation nor endocytosis, related to Figure 6**
(A-B) To exclude that the candidate PD-causing variant affects ENDOA1 conformation, we purified recombinant $\text{ENDOA1}^{\text{WT}}$ and $\text{ENDOA1}^{\text{G276V}}$ and analyzed them by SAXS. The scattering profiles (A) of *Human* $\text{ENDOA1}^{\text{WT}}$ (in the absence and presence of $Ca^{2+}$) and $\text{ENDOA1}^{\text{G276V}}$ are superimposable, excluding large conformational rearrangements in $\text{ENDOA1}^{\text{G276V}}$. The scattering profile of $\text{ENDOA1}^{\text{WT}}$ in the absece of $Ca^{2+}$ shows a small tendency for radiation-induced aggregation under the conditions used. (B) The Kratky plots of *Human* $\text{ENDOA1}^{\text{WT}}$ (in the absence and presence of $Ca^{2+}$), $\text{ENDOA1}^{\text{G276V}}$ in 20 mM MOPS +150 mM NaCl +1 mM DTT +5 % glycerol confirmed that the proteins were well folded.
(C) Westen blot of $endoA^{-/-}$ mutants expressing $SH3GL2^{WT}$ and $SH3GL2^{G276V}$ under a pan-neuronal driver (*nSyb-Gal4*). Blots were probed with anti-ENDOA1 and anti-GAPDH

antibodies. (C') Quantification of ENDOA1 signal intensity relative to GAPDH intensity. Error bars represent mean ± SEM; statistical significance calculated with an ordinary one-way ANOVA with Tukey's multiple comparison test: ns, not significant. Experiment performed in three independent biological replicates.

(D) Representative images of boutons loaded (1 min, 90 mM KCl, 1.5 mM $CaCl_2$) with FM 1-43 (4 µM) and quantification (D') of the following genotypes: control (*nSyb-Gal4/+*), *endoA*$^{-/-}$ animals expressing *SH3GL2*$^{WT}$ and *SH3GL2*$^{G276V}$ under the control of *nSyb-Gal4*. Scale bar: 5 µm. Statistical significance calculated with an ordinary one-way ANOVA with Tukey's multiple comparison test: ns, not significant, n ≥ 7 larvae (28 NMJs) per genotype.

**Supplementary Figure 7 Characterization of iPSC knock-in line and floor plate neural progenitors, related to Figure 7**

(A) CGH array of the edited iPSCs line showing no chromosomal aberrations following gene editing.

(B) representative maximum projection confocal images of control and gene edited (SH3GL2$^{GV}$) iPSCs stained for the pluripotency markers NANOG, OCT4 and SOX2. Scale bar: 150 µm.

(C) Representative maximum projection confocal images of floor plate progenitors (day 16) stained for the indicated floor plate markers to assess the degree of ventralization. Scale bar: 100 µm.

**Supplemental Movie 1 *In vivo* single particle tracking of EndoA, related to Figure 4**

EndoA$^{mEos3.1}$ single molecule imaging at the motor nerve terminal. Movie acquired at 20 Hz. Scale bar: 5 µm.
